# CHORD: a framework for cross-species single-cell integration across gene, cell and cell-type levels

**DOI:** 10.64898/2026.04.19.719426

**Authors:** Yitao Lin, Xunuo Zhu, Xuheng Zhou, Xue Zhang, Guoxin Cai, Wenyi Zhao, Jingqi Zhou, Jie Liu, Qiang Zhu, Menghan Zhang, Binbin Zhou, Xun Gu, Zhan Zhou

**Affiliations:** State Key Laboratory of Advanced Drug Delivery and Release Systems & Innovation Institute for Artificial Intelligence in Medicine, College of Pharmaceutical Sciences, Zhejiang University, Hangzhou, 310058, China; Shanghai Innovation Institute, Shanghai, 200030, China; Zhejiang Provincial Key Laboratory of Anti-Cancer Drug Research & MOE Engineering Research Center of Innovative Anticancer Drugs, Zhejiang University, Hangzhou, 310018, China; School of Public Health, Shanghai Jiao Tong University School of Medicine, Shanghai, 200025, China; Department of Computer Science, City University of Hong Kong, Kowloon, Hong Kong SAR, 999077, China; College of Computer Science and Technology, Zhejiang University, Hangzhou, 310027, China; State Key Laboratory of Genetics and Development of Complex Phenotypes, Center for Evolutionary Biology, Human Phenome Institute, Zhangjiang Fudan International Innovation Center, School of Life Sciences, Fudan University, Shanghai, China; Research Institute of Intelligent Complex Systems, Fudan University, Shanghai, China; School of Computer and Computing Science, Hangzhou City University, Hangzhou, 310015, China; Department of Genetics, Development and Cell Biology, Iowa State University, Ames, IA, 50011, USA

## Abstract

Quantifying cross-species relationships among cell types from single-cell transcriptomic data can reveal both conserved and divergent patterns of cell-type hierarchies. However, existing cross-species integration methods can be limited in modeling genes beyond orthologs by leveraging cell-type-resolved transcriptional context, or in learning explicit type-level representations. Here we present CHORD, a cross-species integration framework that jointly learns representations of genes, cells and cell types. We demonstrate that CHORD can integrate cross-species single-cell atlases and support cell-type annotation with unknown cell-type detection. In the frog–zebrafish embryogenesis and mammalian motor cortex atlases, CHORD infers cell-type trees that place conserved cell types from different species in relative proximity and summarize hierarchical relationships among cell types. CHORD also supports cross-species comparison of continuous phenotypic variation by placing embryonic cells along an aligned developmental timeline. CHORD further yields gene embeddings that capture orthologous and functional relationships, and gene importance scores linking genes to cell types.

## 1 Introduction

Establishing cross-species relationships among cell types at the transcriptomic level is essential for identifying conserved hierarchical organization of cell types and dissecting how it diverges across evolution [1, 2]. Single-cell RNA sequencing (scRNA-seq) has facilitated large-scale cellular atlases within diverse species [3–6], providing a transcriptomic basis for characterizing cell types. However, cell-type annotations are often inconsistent across species, reflecting both genuine divergence in transcriptional programs and methodological differences in sampling and annotation criteria [7–9]. This inconsistency makes systematic cross-species comparison difficult, motivating integration frameworks that align cell types across single-cell atlases in a shared latent space [10].

To integrate single-cell atlases across species, existing approaches mainly face two challenges: how to relate genes across species, and how to infer cell-type relationships. For modeling gene relationships across species, many methods restrict features to one-to-one orthologs [11], which excludes non-orthologous and species-specific genes that potentially help resolve lineage identity and subtype structure [12, 13]. Meanwhile, assuming one-to-one ortholog expression is directly comparable across species can introduce cross-species shifts, since regulatory evolution can substantially alter cell-type-specific expression even for one-to-one orthologs [14]. To broaden gene coverage beyond ortholog-restricted gene sets, recent methods have leveraged protein language models (PLMs) by encoding functional priors derived from amino-acid sequences [15, 16]. However, such sequence-based gene embeddings do not explicitly encode cell-type-resolved transcriptional context from single-cell expression profiles [17]. Such context can be important for relating genes to cell-type expression programs in cross-species integration. For modeling cell-type relationships, most existing cross-species integration methods learn representations at the level of individual cells, and cell-type relationships are typically inferred by comparing cell-level neighborhoods [18]. Cellular identity is inherently hierarchical, which motivates learning explicit representations for cell types to enable direct type-to-type comparison and characterization of hierarchical relationships [19, 20]. Consequently, these considerations highlight the need for cross-species integration frameworks that (i) model genes beyond orthologs while incorporating cell-type-resolved transcriptional context, and (ii) provide cell-type representations that make multi-resolution relationships among cell types explicit.

Here, we present CHORD (Cross-species Hierarchical Orthologous Relationship Discovery), a cross-species integration framework for joint representation learning across gene, cell and cell-type levels. CHORD learns gene embeddings for all input genes by combining one-to-one ortholog anchors with cell-type-averaged expression profiles, and links cells to their types to model explicit cell-type embeddings. We evaluate CHORD on core cross-species tasks, including atlas integration and cell-type annotation with unknown cell-type detection. Using cell-type embeddings of CHORD, we build data-driven cell-type trees in a zebrafish–frog embryogenesis atlas and a human–marmoset–mouse brain atlas. We then show that CHORD is robust to annotation resolution, preserving broad cell-type organization while supporting fine-grained alignment under multi-resolution supervision. We further place embryonic cells along a shared developmental progression to quantify cross-species differences in transcriptional dynamics. Finally, we analyze the gene-level outputs of CHORD, showing that the learned gene embeddings capture orthologous relationships and functional relatedness across species, and that gene importance scores provide an interpretable link between genes and cell types.

## 2 Results

### 2.1 Overview of CHORD

To integrate multi-species scRNA-seq data, we developed CHORD, a representation learning framework that jointly models genes, cells and cell types. CHORD takes as input: (i) scRNA-seq datasets including within-species cell-type annotations and (ii) one-to-one orthologs. Motivated by the need to represent cell types explicitly, CHORD employs a dual-stream architecture, enabling it to model both single-cell variation and cell-type semantics (Fig. 1a). To obtain cell embeddings, CHORD applies random gene masking to the expression profile of each cell, multiplies the masked expression matrix by a learnable gene embedding matrix in which each gene is represented by a vector, and passes the resulting aggregated gene representation through a cell multilayer perceptron (MLP). For cell-type embeddings, CHORD parameterizes a learnable cell-type prototype for each cell type and maps these prototypes through a cell-type MLP. The weights of both the cell MLP and the cell-type MLP are shared across species.

**Fig. 1.**
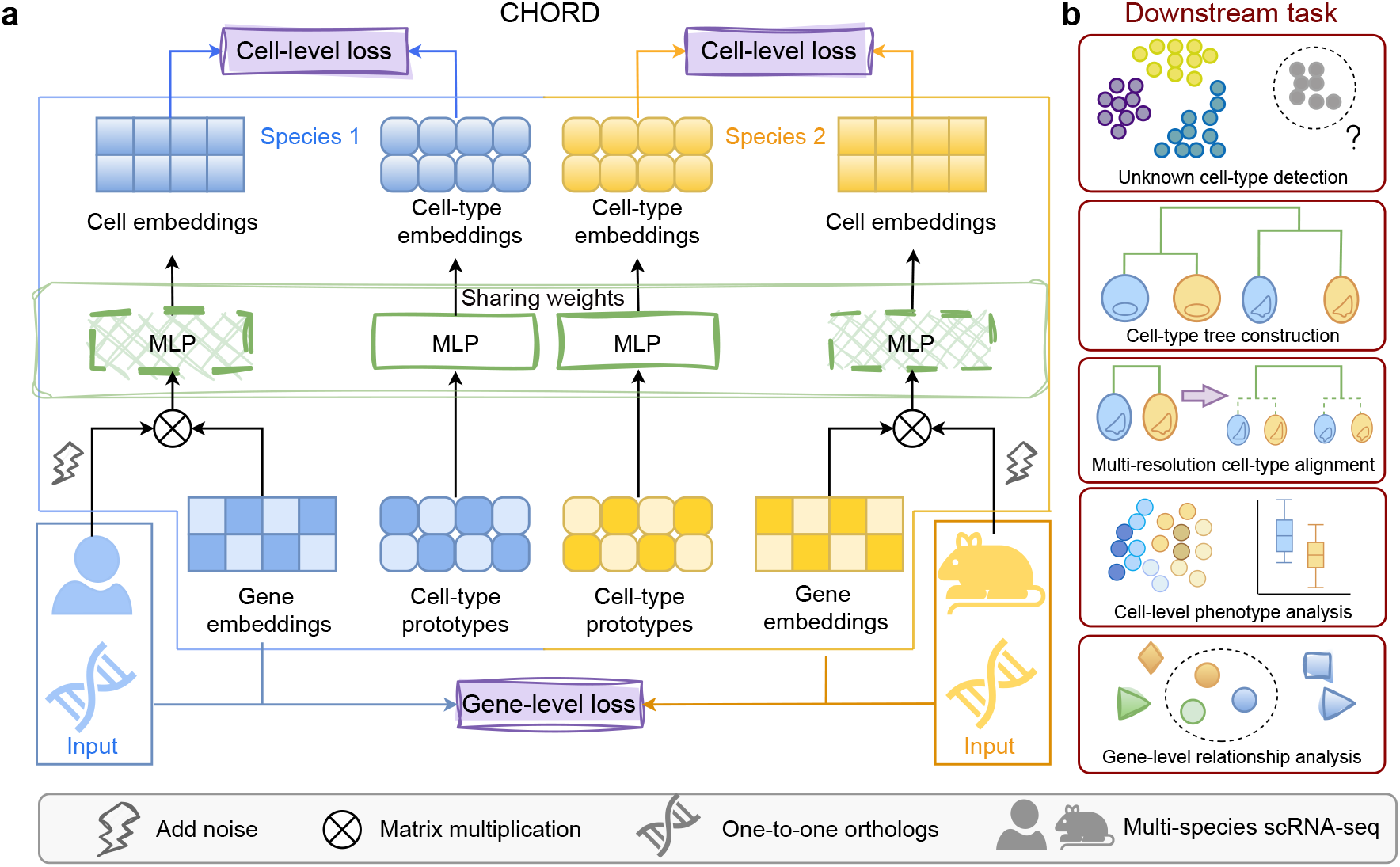
(a) Overview of CHORD for cross-species integration. CHORD supports integration of more than two species. For clarity, the diagram shows the formulation for a single species pair. CHORD takes scRNA-seq expression data with within-species cell-type annotations from each species, together with one-to-one orthologs, as inputs. A dual-stream architecture is applied for each species to learn cell and cell-type embeddings, comprising a learnable gene embedding matrix and a cell-type prototype matrix, together with two MLPs whose weights are shared across species. The gene-level loss is computed for each species pair and summed over all pairs, whereas the cell-level loss is computed within each species. (b) CHORD jointly learns gene, cell, and cell-type embeddings that support diverse downstream analyses across multiple biological levels.

We designed CHORD with two complementary objectives: a gene-level loss and a cell-level loss. The gene-level loss aligns gene embeddings across species for all input genes. By leveraging cell-type-averaged expression profiles together with one-to-one ortholog anchors, CHORD propagates cross-species information from orthologs to other input genes and guides gene embeddings to capture cell-type-level transcriptional context (Methods). The cell-level loss is computed within each species to preserve cell-type separability. CHORD leverages a prototype-based contrastive objective that pushes cell embeddings toward their corresponding cell-type embeddings while repelling them from other cell-type embeddings, thereby allowing cell-type embeddings to act as a compact summary of their member cells. A two-stage training schedule is used, in which we first optimize the gene-level loss alone and then jointly optimize both losses. CHORD supports diverse downstream applications across multiple representation levels. In our study, cell-type embeddings were mainly used for cell-type tree construction, unknown cell-type detection, and multi-resolution integration, whereas cell embeddings were used for phenotype analysis and gene embeddings for gene relationship analysis (Fig. 1b).

### 2.2 Benchmarking cross-species integration, cell-type annotation and unknown-cell separation

We designed benchmarks to test whether CHORD can (i) achieve high-quality cross-species integration, (ii) annotate cells while detecting unknown cell types within species and across species, and (iii) separate withheld cell types among cells predicted as unknown. We used two publicly available cross-species scRNA-seq datasets, including an evolutionarily distant pair (frog–zebrafish embryogenesis; total *n* = 160,306) [21] and a three-species atlas (human–mouse–marmoset primary motor cortex; total *n* = 305,550) [22]. We compared CHORD against representative baselines spanning different levels of cell-type label supervision: (i) fully unsupervised models that do not use cell-type labels (scVI [23]); (ii) supervised methods that use within-species labels without harmonized cross-species labels (SATURN [16]), the same supervision level as CHORD; and (iii) supervised methods that can exploit harmonized cross-species labels (scANVI [24], scGen [25]).

To quantify integration performance, we evaluated two aspects: biological conservation, assessed by whether annotated cell types remained well separated and consistently recovered after integration, and batch correction, reflecting the extent to which species-driven batch effects are mitigated in the integrated embedding (Methods). CHORD achieved better biological conservation and batch correction scores on both datasets (Fig. 2a and Supplementary Fig. 1). For biological conservation, CHORD improved over the next best-performing method (SATURN) by 6.08% on the evolutionarily distant frog–zebrafish pair. For batch correction, CHORD slightly exceeded scANVI on both datasets, with relative improvements of 2.62% (frog–zebrafish) and 1.30% (brain), suggesting that CHORD can achieve strong integration without relying on harmonized cross-species labels.

**Fig. 2.**
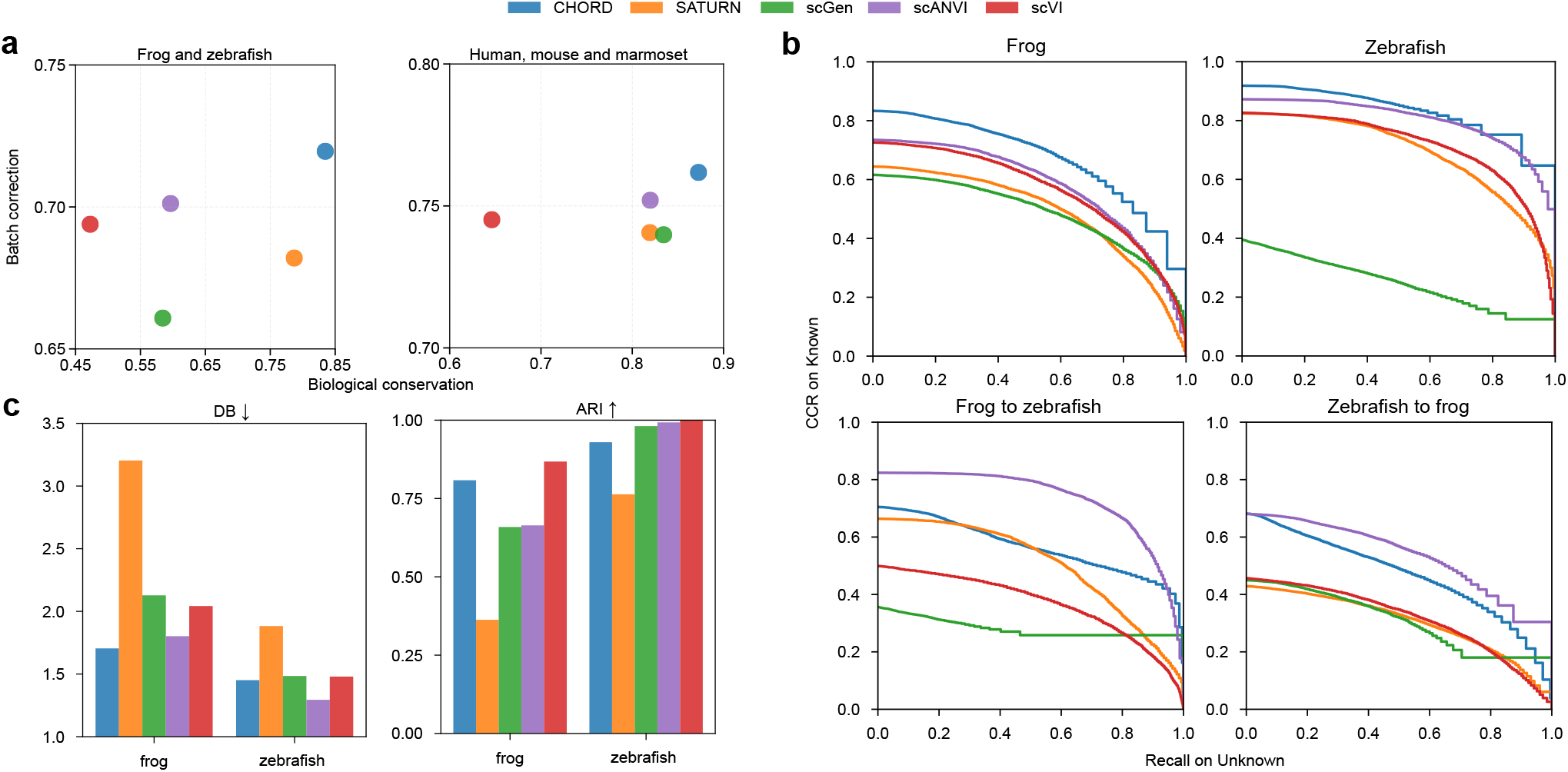
Benchmarking cross-species integration, cell-type annotation and unknown-cell separation. (a) Integration performance of CHORD compared to baselines. The biological conservation score is the average of cell-type NMI, cell-type ARI, cell-type ASW and trajectory score (only applicable in the frog–zebrafish datasets). The batch correction score is the average of ASW batch score and graph connectivity score, where species is taken as the batch variable. (b) Cell-type annotation with unknown cell-type detection. The top panels show within-species annotation performance, where training and test cells come from the same species. The bottom panels show cross-species label transfer, where test cells come from a different species than the training data. In each panel, the correct classification rate of known types (CCR, y-axis) counts a known cell as correct only if it is not detected as unknown and its predicted label matches the ground truth, while recall of unknown cell types (x-axis) measures the fraction of unknown cells that are detected (rejected) as unknown. (c) Clustering quality of withheld cell types. The left panel reports the Davies–Bouldin (DB) index (lower is better), and the right panel reports the adjusted Rand index (ARI; higher is better).

Next, we evaluated CHORD for both cell-type annotation and unknown cell-type detection in within-species classification and cross-species label transfer. Using the frog–zebrafish embryogenesis dataset, the test set consisted of two components: held-out cells from known cell types and all cells from five withheld cell types in each species (Supplementary Note 4). We measured the accuracy of annotating known cell types and the ability to detect (reject) unknown cell types using two metrics: the correct classification rate (CCR) for known cell types and the recall of unknown cell types. All methods exhibited a trade-off: CCR on known types typically decreases as recall on unknown types increases (Fig. 2b). In within-species classification, CHORD achieved higher CCR at matched unknown-type recall across most of the curve. At an unknown-type recall of 0.7, CHORD achieved a CCR of 0.623 in frog, compared with 0.519 for scANVI, 0.502 for scVI, 0.434 for SATURN and 0.430 for scGen. A similar pattern was observed in zebrafish, where CHORD remained competitive at the same recall level (CCR = 0.804), comparable to scANVI (CCR = 0.784) and higher than the other baselines. The same trend held for cross-species label transfer, where CHORD ranked second only to scANVI across much of the curve. At an unknown-type recall of 0.7, CHORD achieved a CCR of 0.399 in zebrafish-to-frog transfer and 0.504 in frog-to-zebrafish transfer, outperforming scVI, SATURN and scGen in both directions.

Following unknown cell-type detection, we examined whether cells from the withheld cell types formed distinct clusters in the integrated embedding. We used four metrics to assess both intrinsic cluster quality (Davies–Bouldin index, DB, and average silhouette width, ASW) and consistency with reference cell-type labels (adjusted Rand index, ARI, and normalized mutual information, NMI). No single method performed best across all clustering metrics in both species (Fig. 2c and Supplementary Fig. 2). Across the 8 metric–species comparisons, CHORD was among the top two methods in 5 cases. In frog, it achieved the highest ASW and the lowest DB. In zebrafish, CHORD ranked second in DB, whereas scANVI, scVI and scGen performed better in ARI, NMI and ASW. We further visualized the embeddings of cells from the withheld cell types using Uniform Manifold Approximation and Projection (UMAP) [26]. Consistent with the clustering metrics, CHORD produced compact and well-separated clusters from the withheld cell types in frog, whereas SATURN and scGen showed greater mixing across cell types and scANVI more often split the same withheld type into multiple clusters (Supplementary Fig. 3). In zebrafish, CHORD retained partial separation among the withheld cell types (Supplementary Fig. 4).

### 2.3 CHORD integrates cell atlases and constructs cell-type trees across species

To examine the cross-species organization learned by CHORD at both the single-cell and cell-type levels, we applied CHORD to the frog–zebrafish embryogenesis dataset and the primary motor cortex atlas from human, marmoset, and mouse. Visualizing the integrated cell embeddings with UMAP, we observed that cells from different species largely intermixed within the same annotated cell types, while different cell types remained separated (Fig. 3a and Supplementary Fig. 6a). Beyond visualizing cell embeddings with UMAP, we constructed cell-type trees using cell-type embeddings learned by CHORD to characterize relationships among annotated cell types (Methods). In the inferred cell-type trees, conserved cell types from different species were positioned in relative proximity, and cell types formed coherent hierarchies (Fig. 3b and Supplementary Fig. 6b).

**Fig. 3.**
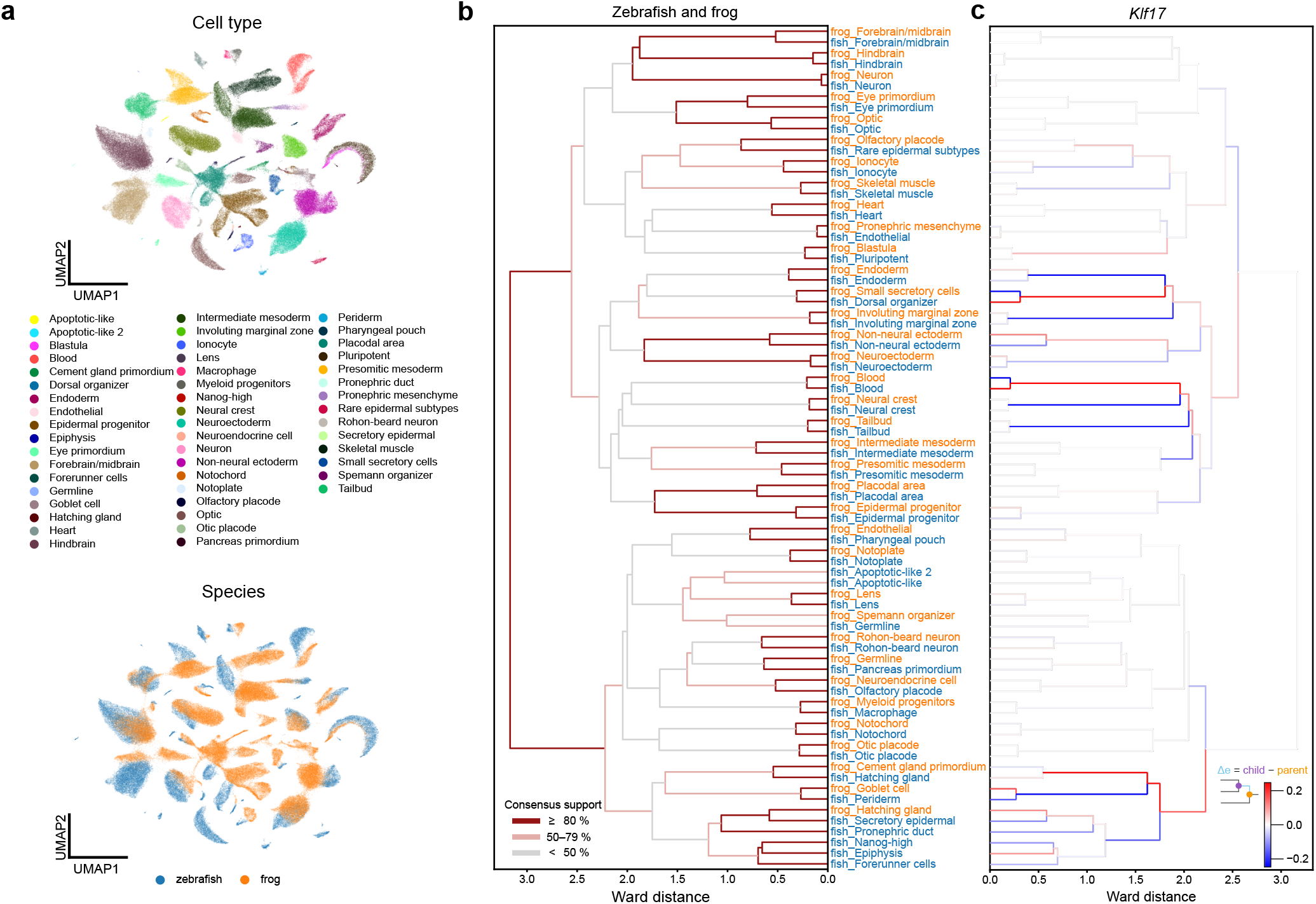
CHORD integrates frog and zebrafish datasets to learn a cross-species cell embedding space and construct a cell-type tree. (a) UMAP visualization of joint cell embeddings across zebrafish (63371 cells) and frog (96935 cells), integrated by CHORD. Colors denote cell-type annotations (top) and species annotations (bottom). (b) Cross-species cell-type tree inferred from CHORD cell-type embeddings using hierarchical clustering (Ward linkage). The x-axis shows Ward distance, where smaller distances indicate higher similarity. Leaves on the y-axis correspond to annotated species-specific cell types. Branch color indicates fuzzy consensus support across epoch-wise trees inferred from the final training epochs (low < 0.5, intermediate 0.5–0.8, high ⩾0.8). Higher support indicates internal splits that are consistently recovered across epochs. (c) Branch-wise changes in *Klf17* ortholog expression mapped onto the cell-type tree in (b) (same topology, mirrored for visualization). For each leaf cell type, we computed the average normalized log-transformed expression across cells; internal nodes are assigned the unweighted mean of their descendant leaf cell-type averages. The expression change along each branch is defined as the difference between a node and its immediate ancestor (child minus parent). Red indicates increased expression and blue indicates decreased expression along the corresponding branch.

We examined whether the cell-type tree captures broader-scale relationships among cell-type groups. For the frog–zebrafish embryogenesis dataset, neuron, hindbrain, and forebrain/midbrain cell types were not only aligned across zebrafish and frog but also co-clustered with high consensus support, consistent with a canonical central nervous system (CNS) organization. Similarly, for the primary motor cortex dataset, vascular-associated classes, including pericytes, vascular and leptomeningeal cells (VLMCs), and endothelial cells, co-clustered with high consensus support, consistent with their molecular characterization as major blood vascular and vessel-associated cell types in the brain [27] (Supplementary Fig. 6b). We next assessed whether the tree captures relative transcriptomic similarity among related cell-type groups. Within the CNS branch, the Ward distance between matched hindbrain cell types across species was smaller than that for forebrain/midbrain (smaller distance indicates higher similarity), in line with a prior report that hindbrain patterning programs are conserved across vertebrates [28].

To provide a transcriptional interpretation of the cell-type trees, we mapped ortholog expression onto them by computing mean expression for each leaf cell type and quantifying parent–child changes along internal splits (Supplementary Note 6). In the embryogenesis tree, zebrafish hatching gland (HG) was placed closer to the frog cement gland (CG) primordium than to the frog HG. From a spatiotemporal perspective, zebrafish HG and the frog CG primordium were both anterior, secretory structures that were identifiable early in development [29, 30], whereas frog HG cells coalesced into a characteristic dorsoanterior head pattern only during neurulation [31]. At the molecular level, along the branch leading to the zebrafish HG and the frog CG primordium, *Klf17*, a conserved regulator of early surface ectoderm programs [32, 33], showed a continuous increase (Fig. 3c). In contrast, along the branch containing the secretory epidermis in zebrafish and the frog HG, *Klf17* showed a weaker or opposite change. Similar branch-wise increases were observed for *Xbp1*, a regulator of endoplasmic reticulum (ER) stress and the unfolded protein response [34], and *Cd63*, a tetraspanin involved in endosomal trafficking and secretory vesicle/exosome pathways [35] (Supplementary Fig. 5). Together, these spatiotemporal and expression patterns were consistent with functional similarity between zebrafish HG and the frog CG primordium. In the mammalian motor cortex tree, human VLMCs were placed closer to pericytes than to non-human VLMCs. This placement may reflect annotation or compositional differences across datasets. Considering that the human datasets lacked a dedicated pericyte annotation, and given the known shared mesenchymal lineage and transcriptomic continuum between these cell types, we hypothesized that the cells labeled as human VLMC may include pericyte-like or transitional states. We observed upregulation of *CYP1B1* along the branch ending at the human VLMC leaf (Supplementary Fig. 6c). *CYP1B1* had been reported to be constitutively expressed in pericytes [36], in line with a pericyte-like component within the human VLMC annotation.

### 2.4 Robustness to annotation resolution via hierarchical fine-tuning

We assessed the sensitivity of CHORD to annotation resolution by training it with both low- and high-resolution cell-type annotations. These annotation resolutions correspond biologically to broader cell-type groups and fine-grained distinctions within them. We used a hierarchical fine-tuning strategy in which CHORD was first trained with low-resolution annotations and then fine-tuned on high-resolution annotations by introducing a new set of high-resolution cell-type prototypes and optimizing a cell-level loss (Fig. 4a and Supplementary Note 7). We compared three training settings in the primary motor cortex dataset: low-resolution-only, high-resolution-only, and low-to-high (hierarchical fine-tuning).

**Fig. 4.**
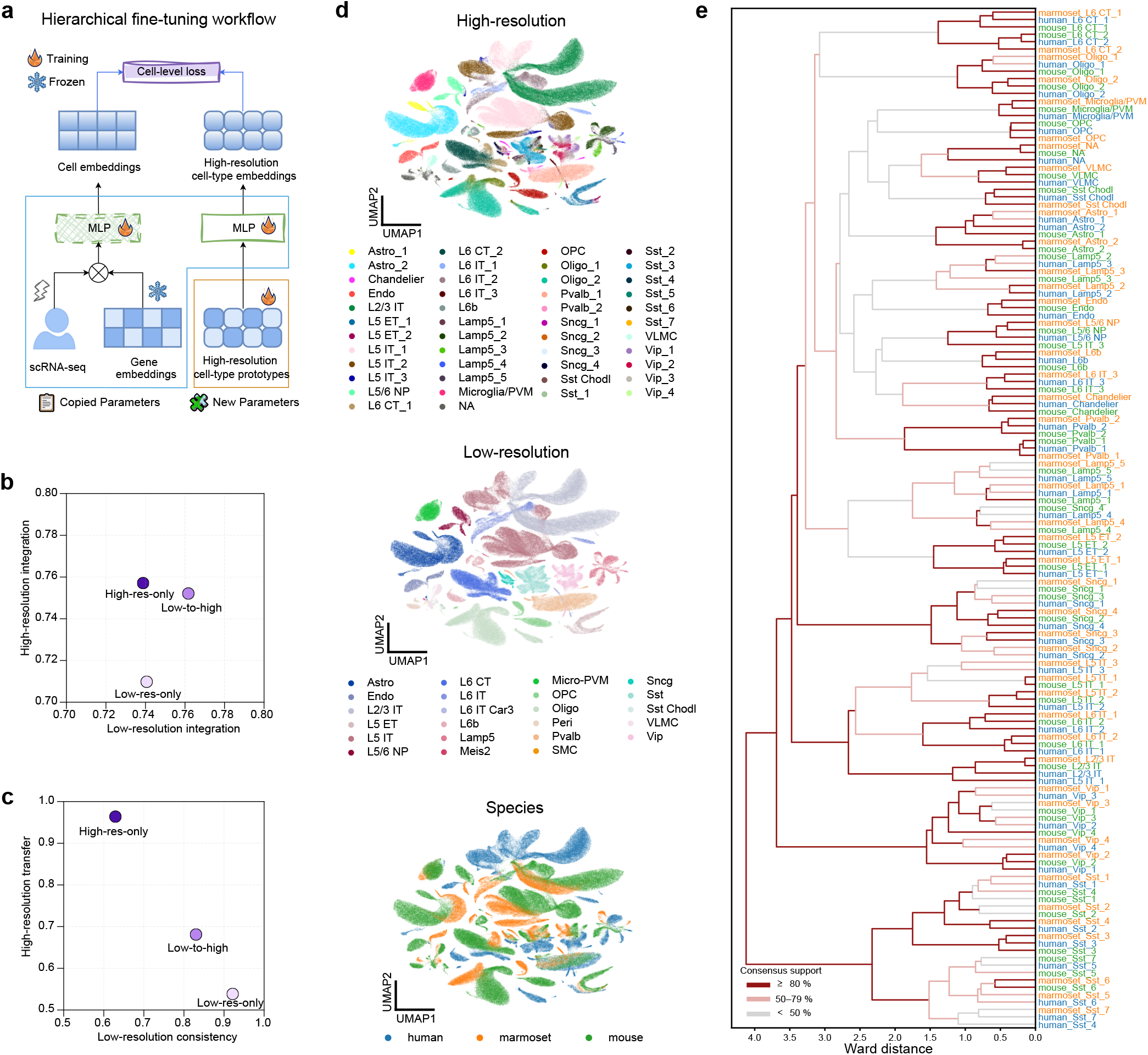
Evaluation of CHORD robustness to annotation resolution via hierarchical fine-tuning. (a) Hierarchical fine-tuning strategy. CHORD is first trained with low-resolution labels. During fine-tuning, a new set of high-resolution cell-type prototypes is introduced to represent high-resolution cell types. Gene embeddings are frozen, while the MLPs in both branches are initialized from the low-resolution-trained model and remain trainable. The model is optimized with a cell-level loss on high-resolution labels. Fire icons denote trainable parameters; snowflake icons denote frozen parameters. (b) Comparison of integration performance at low and high annotation levels for low-resolution-only, high-resolution-only, and low-to-high models. Axes show overall integration scores at the low and high annotation levels, computed as a weighted sum of the biological conservation score (60%) and the batch correction score (40%). (c) Performance of the three models on hierarchical consistency and cross-species transfer of high-resolution labels. Hierarchical consistency is evaluated by ARI and NMI between low-resolution labels induced from the high-resolution cell-type tree and the low-resolution evaluation groups, whereas cross-species label transfer is evaluated at the high level using accuracy and macro F1. For high-resolution evaluation under the low-resolution-only setting, we introduce high-resolution cell-type prototypes and update the prototypes only to obtain high-resolution cell-type embeddings. (d) UMAP visualization of CHORD cell embeddings under the hierarchical fine-tuning strategy, with cells colored by high-resolution labels, low-resolution labels, and species. (e) A cell-type tree inferred from the low-to-high model. Leaf labels denote species and high-resolution cell types.

We first assessed integration performance under both low- and high-resolution annotations, using an overall integration score defined as a weighted average of the biological conservation and batch correction scores. Hierarchical fine-tuning achieved intermediate integration performance at the high annotation level between the low-resolution-only and high-resolution-only models (Fig. 4b). Notably, batch correction scores at the low annotation level were slightly higher than those of the other two models, suggesting that introducing high-resolution supervision can preserve and even improve integration quality at the low annotation level (Supplementary Fig. 8). Next, we evaluated the structural validity of the learned cell-type organization using two tests: (i) hierarchical consistency of the low-resolution organization, to assess whether broad cell-type organization was maintained, and (ii) cross-species transfer of high-resolution labels, to assess whether fine-grained cell types were aligned across species. The low-to-high model showed intermediate performance on both tests, lying between the low-resolution-only and high-resolution-only models (Fig. 4c and Supplementary Fig. 9). Consistent with these quantitative trends, UMAP of cell embeddings from the low-to-high model showed separation of broad cell-type groups, alongside cross-species co-localization of high-resolution cell types within each group (Fig. 4d). In the cell-type trees, the low-resolution-only model preserved broad cell-type organization but did not capture fine-grained cross-species alignment for high-resolution cell types such as Sncg interneurons (Supplementary Fig. 7a). Conversely, the high-resolution-only model better aligned fine-grained cell types across species but struggled to preserve structure in broad cell-type groups, such as L6 corticothalamic (L6 CT) neurons (Supplementary Fig. 7b). By integrating supervision from both resolutions, the low-to-high model maintained broad cell-type organization while achieving fine-grained cross-species alignment among high-resolution cell types (Fig. 4e).

### 2.5 CHORD enables cross-species developmental alignment and transcriptional tempo analysis

To illustrate how CHORD cell embeddings can be linked to continuous phenotypes, we used embryonic developmental progress as a test case by aligning frog stages to zebrafish time points and quantifying transcriptomic change along the aligned timeline. To align different time labels across species, we computed a time–similarity matrix using symmetric 1-nearest-neighbor (1-NN) cosine distances between the sets of CHORD cell embeddings from each frog stage and each zebrafish time point (hpf) (Supplementary Note 8). From this matrix, we obtained an initial, discrete alignment by assigning each frog stage to the most similar zebrafish time point (Fig. 5a). Frog Stages 16–22 were all mapped to the same zebrafish endpoint (24 hpf) and thus provided little additional time resolution beyond Stage 16. We therefore retained frog Stage 16 as the boundary but excluded stages after Stage 16 from subsequent developmental analysis. Within the retained window (frog Stages 8–16), to avoid many-to-one mappings that create flat segments, we linearly interpolated the mapped times. As zebrafish time increased, the mapped frog stage rose with a decreasing slope, indicating that later increases in zebrafish time correspond to progressively smaller stage advances in the frog (Supplementary Fig. 10a).

**Fig. 5.**
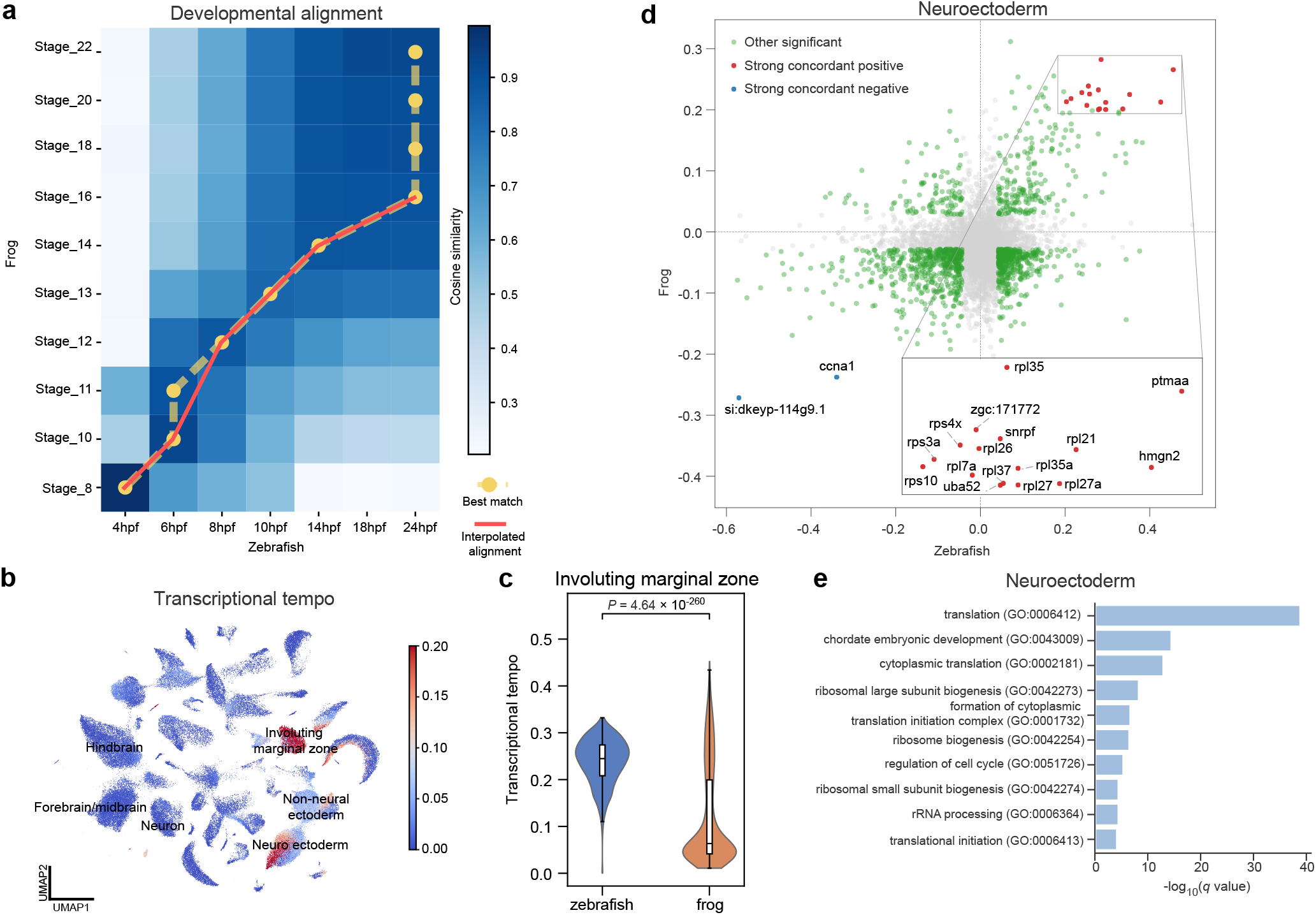
CHORD aligns frog and zebrafish developmental time, quantifies transcriptional tempo, and links tempo-associated genes to molecular programs. (a) Developmental alignment between frog stages and zebrafish hours post-fertilization (hpf) using the CHORD cell embeddings. The heatmap shows time–similarity computed from symmetric 1-nearest-neighbor (1-NN) cosine distances between frog stages and zebrafish time points. Yellow circles mark the best-match zebrafish time for each frog stage. The red line interpolates these assignments within the retained window (frog Stages 8–16) to obtain a continuously increasing stage-to-hpf mapping. Along this mapping, frog Stage 11 is assigned 7 hpf. (b) Transcriptional tempo estimated for individual cells using the CHORD cell embeddings and visualized on a UMAP. The color scale is truncated at 0.2 to prevent a few extreme values from dominating the display. (c) Distribution of transcriptional tempo within the involuting marginal zone (IMZ). *P* values are computed using a two-sided Wilcoxon rank-sum test. (d) Cross-species associations between gene expression and transcriptional tempo. Each point is a gene matched by one-to-one ortholog, positioned by its Spearman correlation in zebrafish (x-axis) and frog (y-axis). Red denotes strong concordant positive correlations in both species (Spearman *ρ* ⩾ 0.2 and *q* < 0.05 in each species); blue denotes strong concordant negative correlations (Spearman *ρ* ⩽ −0.2 and *q* < 0.05 in each species); green denotes other cross-species significant genes (*q* < 0.05 in each species) after excluding the red/blue sets. Each *q* value was obtained by applying the Benjamini–Hochberg procedure to the per-gene correlation *P* values within each species. (e) Gene-set enrichment analysis of the broader cross-species positive set (Spearman *ρ* > 0 and *q* < 0.05 in each species) was performed using Gene Ontology Biological Process terms, and enrichment *P* values were adjusted by the Benjamini–Hochberg procedure.

To quantify how quickly the transcriptomic state of a cell shifts along the aligned developmental timeline, we defined an index termed transcriptional tempo. For each cell, transcriptional tempo was computed as the cosine distance between its CHORD embedding and that of its nearest neighbor at the subsequent aligned time point within the same species, normalized by the corresponding time interval. Transcriptional tempo varied across cell types. Cells from the involuting marginal zone (IMZ) exhibited the highest transcriptional tempo, with elevated values also observed in subsets of neuroectoderm and non-neural ectoderm (Fig. 5b). We next compared the transcriptional tempo between species on the aligned timeline. In the IMZ, zebrafish exhibited higher transcriptional tempo than the frog (*P* = 4.64 × 10^−260^; Fig. 5c). In the forebrain/midbrain, transcriptional tempo values were close to zero in both species, showing a much smaller cross-species difference than in the IMZ (Supplementary Fig. 10b). To connect transcriptional tempo differences to molecular programs, we calculated the Spearman correlation between ortholog expression and transcriptional tempo within neuroectoderm in each species. Among the genes highlighted as strong concordant positives, many were ribosomal protein genes (RPL/RPS) (Fig. 5d). We performed gene-set enrichment analysis using a broader set of genes with concordant positive correlations across species (Spearman *ρ* > 0 and *q* < 0.05 in each species). The top enriched terms included translation and chordate embryonic development (Fig. 5e), consistent with increased protein-synthesis programs during development [37].

### 2.6 Aligned gene embeddings discover biologically meaningful cross-species relationships

CHORD produced gene embeddings across species in a shared space, allowing genes to be compared by cosine similarity. By visualizing the learned gene embeddings with UMAP, we qualitatively observed both cross-species overlap and species-specific separation (Supplementary Fig. 11a). Next, we evaluated the gene-level outputs of CHORD from three complementary aspects: (i) alignment of one-to-one ortholog pairs in the gene embeddings, (ii) cross-species functional consistency of embedding neighborhoods, and (iii) gene importance scores and the resulting cell-type gene signatures across species (Supplementary Note 9).

To assess whether the gene embeddings capture orthologous relationships, we compared cosine similarities of ortholog pairs to an equal-sized set of randomly paired non-ortholog cross-species genes. Ortholog pairs were significantly shifted toward higher similarity than random pairs (Fig. 6a and Supplementary Fig. 11b). We next used the distribution of ortholog-pair similarities to characterize cross-species divergence in the embedding. Ortholog similarities were highest for human–marmoset and lower for comparisons involving mouse (Fig. 6b), consistent with greater divergence between primates and rodents. To evaluate whether the embedding captures functional neighborhoods within each species and consistent functional signals across species, we tested enrichment of known interactors among each gene’s nearest embedding neighbors using curated protein–protein interaction (PPI) networks in each species, and then combined species-specific *P* values using Fisher’s method. *ERMN*, a myelin gene associated with oligodendrocyte [38], was significant in human (*q*_human_ = 9.5 × 10^−4^) (Supplementary Fig. 11c). Aggregating per-species evidence by meta-analysis, we observed cross-species consistency even when within-species enrichment signals are weak after Benjamini–Hochberg correction (Supplementary Table 2). *AEBP1* showed significant meta-analysis support (*q*_meta_ = 4.1 × 10^−4^). *COL1A2* appeared among the nearest embedding neighbors of *AEBP1* in both human and mouse, and the *AEBP1* –*COL1A2* interaction was independently reported in curated PPI networks for each species (Fig. 6c). To obtain an interpretable gene-level attribution for each cell type, we computed gene importance scores by multiplying input expression by gradients of the cell-type objective, and ranked genes by z-scored importance within each species. In Pvalb interneurons, *ERBB4* and *NXPH1* showed shared high importance scores across species (Fig. 6d), consistent with prior reports implicating *ERBB4* in parvalbumin interneuron function [39] and *NXPH1* in inhibitory synaptic transmission and plasticity [40]. Similarly, in the L6 IT subclass, cadherin-family genes (e.g., *CDH9* /*CDH13*) were highly ranked by importance scores in multiple species (Supplementary Fig. 11d), suggesting that cadherin-associated features were consistently highlighted in this subclass.

**Fig. 6.**
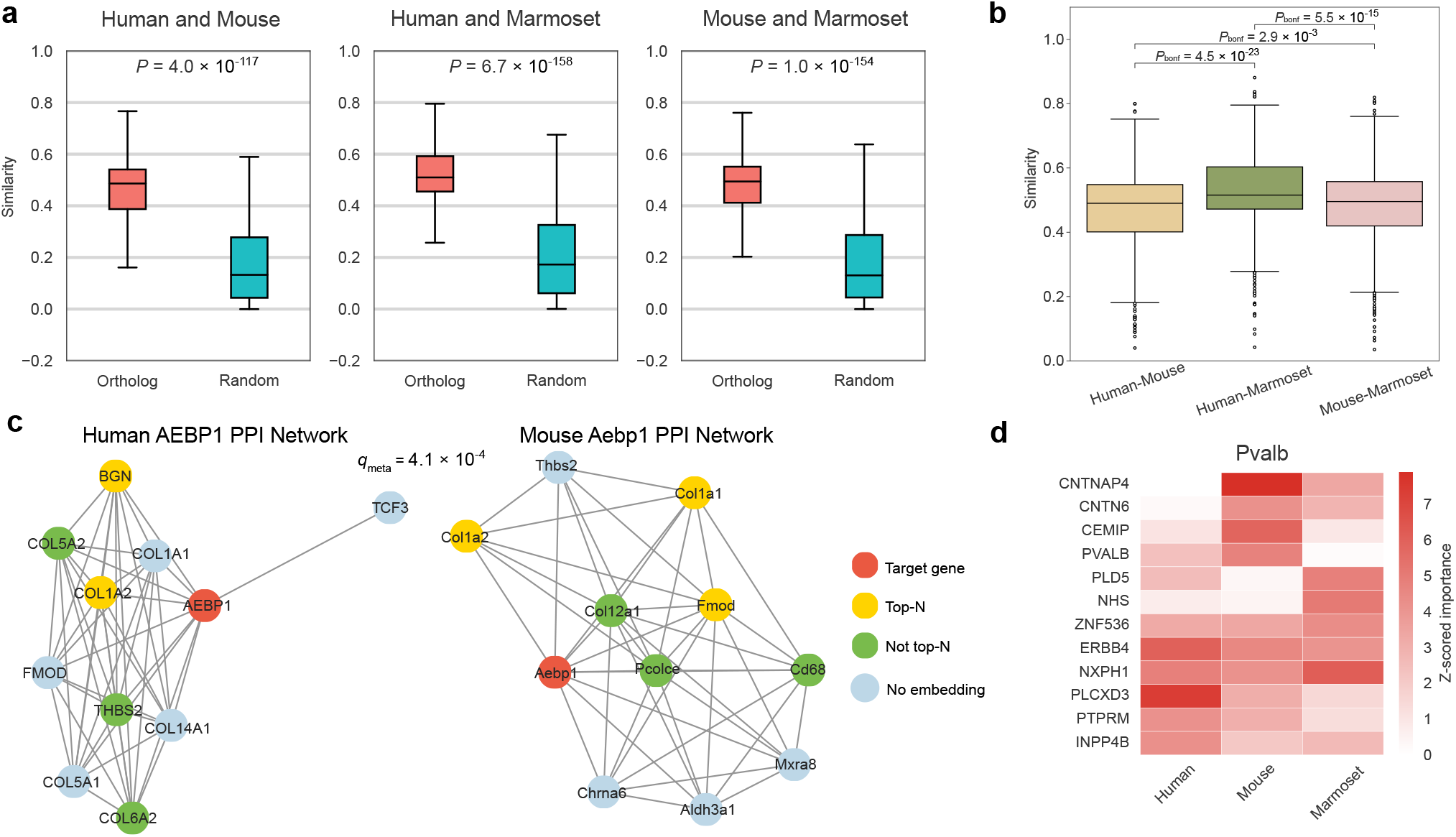
CHORD learns aligned cross-species gene embeddings that capture biological relationships. (a) Distribution of cosine similarities in the gene embeddings for one-to-one ortholog pairs and an equal-sized set of randomly paired non-ortholog pairs. A Wilcoxon rank-sum test was used with the hypothesis that ortholog pairs exhibit higher similarity than random pairs. (b) Distribution of cosine similarity in the embedding for each species pair, computed on one-to-one ortholog triplets shared by human, mouse, and marmoset (*n* = 439 triplets). Pairwise differences were tested using a two-sided Wilcoxon signed-rank test (paired by triplets), with Bonferroni correction for the three comparisons. (c) PPI networks for orthologs of *AEBP1* in human and mouse. Target gene (red); top *N* the nearest embedding neighbors (yellow); other genes with embeddings (green); genes without learned embeddings (blue). *q*_meta_ denotes the meta-analysis *q* value computed by applying the Benjamini–Hochberg procedure to the Fisher-combined enrichment *P* values from human and mouse. (d) Gene importance scores for the Pvalb cell type across species (gradient×input). Heatmap shows the top *N* = 5 genes for each species (Human, Mouse, Marmoset), ranked by z-scored gene importance scores (within-species normalization).

## 3 Discussion

We developed CHORD for cross-species integration by unifying gene-, cell- and cell-type-level modeling within a single framework. CHORD extends cross-species gene representation beyond strict one-to-one orthologs, allowing gene relationships to be inferred from transcriptional context. By explicitly modeling cell-type embeddings, CHORD enables multi-resolution relationships among cell types to be examined across species. Together, these representations expand cross-species analysis beyond cell integration alone, enabling downstream applications at the gene, cell and cell-type levels.

Across two datasets spanning both two- and three-species settings, CHORD achieved strong cross-species integration. We further evaluated CHORD in a more practical annotation setting that jointly considers cell-type annotation and detection of unknown cell types, as novel or incompletely matched cell types are common in cross-species comparisons. CHORD relies only on within-species labels, and its strong performance in this setting highlights its practical utility in comparative analyses where cross-species labels are incomplete or inconsistently defined.

At the cell-type level, CHORD learns cell-type embeddings that support construction of a data-driven cell-type tree. Conventional transcriptomic similarity trees often compute pairwise distances (e.g., correlation- or Euclidean-based) on normalized log-scale expression vectors, and can therefore be sensitive to normalization choices [41] and the cross-species gene set (e.g., ortholog filtering) used for comparison [42]. Instead of deriving the tree directly from normalized expression vectors, CHORD constructs it from learned cell-type embeddings in a shared cross-species latent space, yielding a transcriptome-level summary of cell-type relationships across species. As a complement to UMAP visualization of cell embeddings, the cell-type tree provides a quantifiable view of broader-scale relationships among cell-type groups beyond local neighborhoods. In addition, mapping ortholog expression onto the tree provides a transcriptional interpretation of its topology by showing how gene expression programs change across related cell types. Unlike species phylogeny, which are largely static reference frameworks grounded in prior knowledge and phylogenetic evidence, the CHORD cell-type tree is data-dependent and can vary with dataset composition and annotation conventions across species [43].

At the cell level, the cell embeddings learned by CHORD enable aligned cells to be linked to continuous phenotypes across species. Using developmental progression as an example, we inferred cross-species temporal correspondence from shared cell embeddings and defined transcriptional tempo as a local rate of transcriptomic change along the aligned timeline. The resulting tempo patterns and their associated transcriptional programs illustrate how integrated cell embeddings can support biologically interpretable phenotype-linked analyses.

At the gene level, CHORD provides a shared gene embedding space that extends cross-species gene comparison beyond strict one-to-one orthologs. This representation supports comparative analyses of orthologous and functionally related genes. Although these embeddings are informed by one-to-one ortholog anchors, they are also shaped by transcriptional context and should be interpreted as transcriptomic correspondence rather than direct evidence of homology. In addition, CHORD yields gene importance scores that provide an interpretable means to prioritize cell-type-associated genes and compare these signatures across species.

Like other integration methods, CHORD has several limitations. First, cell-type embedding distances provide a relative transcriptomic similarity metric rather than an absolute, directly interpretable distance scale. When interpreting the cell-type tree with respect to a fixed reference topology (e.g., an ontology or species phylogeny), distance calibration would be required [44, 45]. In addition, broader splits in the inferred cell-type tree can be less stable because hierarchical clustering compresses inter-cluster differences as broad cell-type groups are formed. A possible extension would be to introduce branch-level representations with top-down consistency constraints. Third, because the optimization objective relies on evaluating all species pairs, the computational cost grows quadratically with the number of species. This cost could be alleviated by sparse pairing strategies, such as using a reference species or a species-neighborhood graph. In future work, we will incorporate biological priors (e.g., ontology and phylogeny structure, gene regulatory and interaction knowledge) to extend transcriptome-only alignment toward multi-layer biological representations, enabling more reliable discovery of conserved and divergent cell-type relationships at scale.

## 4 Methods

### Input of CHORD

CHORD takes two inputs. Let Ω = {1, …, *m*} denote the set of species. The first is a set of annotated, preprocessed single-cell RNA-seq datasets from multiple species {**X**′^(1)^, **X**′^(2)^, …, **X**′^(*m*)^}, where the cell-type annotations are provided within each species, without requiring harmonized cross-species labels. For species *i*, we ensure that library-size normalization and log-transformation (log1p) have been applied and select highly variable genes (HVGs) to construct the input matrix 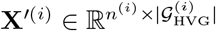, where *n*^(*i*)^ denotes the number of cells and 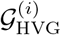 denotes the set of HVGs. The second input consists of one-to-one orthologs between species, which are used to define anchors for modeling cross-species gene relationships.

### Modeling at the Single-Cell and Cell-Type Levels

For each species, CHORD adopts a dual-stream architecture to model both within-species single-cell variation and cell-type semantics in a shared latent space. To calculate cell embeddings, a learnable gene embedding matrix 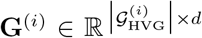 is defined, where *d* is the dimension of the gene embedding space. Each row in **G**^(*i*)^ represents a single gene for species *i*. For species *i*, we apply a stochastic gene expression mask to the preprocessed single-cell gene expression matrix **X**^′(*i*)^, yielding the masked input 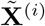 (Supplementary Note 10). The cell embedding 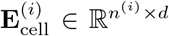 is computed as an expression-weighted linear combination of gene embeddings **G**^(*i*)^, followed by a species-shared cell multi-layer perceptron (MLP) to extract complex features:

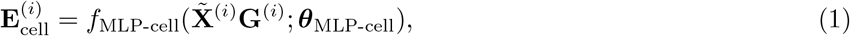

where ***θ***_MLP-cell_ denotes the learnable parameters of the cell MLP.

To represent annotated cell types explicitly, we introduce a learnable, species-specific cell-type prototype matrix 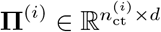, where 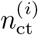 is the number of cell types present in species *i*. Each row in **Π**^(*i*)^ represents a cell-type prototype for species *i*. The cell-type embedding 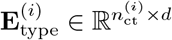 is produced by another cell-type MLP shared across species:

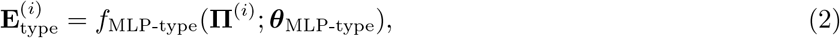

where ***θ***_MLP-type_ denotes the learnable parameters of the cell-type MLP.

### Gene-level loss

We design a gene-level loss to learn aligned gene embeddings that capture cross-species transcriptional associations. For each species pair (*i, j*), we first identify a set of anchor orthologs by restricting one-to-one orthologs to those that are highly variable in both species. We calculate a cell-type-averaged expression matrix 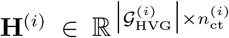, where 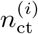 denotes the number of cell types for species *i*. Using the cell-type average expression profiles of these anchor orthologs, we computed pairwise cosine similarities between cell types across species. We then retained only mutual top-*k* nearest-neighbor pairs to define a robust cross-species cell-type correspondence, represented by a matrix 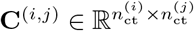 (Supplementary Note 11). These correspondences are then used to define a gene-level supervision signal, encouraging genes to align across species when they exhibit similar expression patterns in corresponding cell-type contexts:

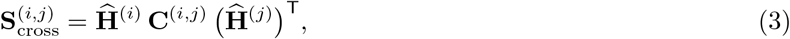

where 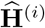 and 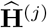 denote row-wise L2-normalized cell-type average expression matrices. The matrix 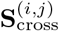 elements are bounded in [−1, 1], which stabilizes optimization (Supplementary Note 12).

Let 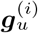 denote the embedding of gene *u* in species *i*, corresponding to the rows of the gene embedding matrix **G**^(*i*)^. We then computed the cross-species gene similarity matrix 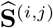 by taking pairwise cosine similarities between gene embeddings from the two species:

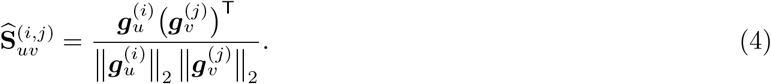

We defined the gene-level alignment loss for species pair (*i, j*) as the mean squared error between the embedding-derived similarity matrix and the association target:

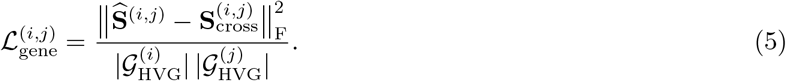

Because this loss is symmetric with respect to swapping *i* and *j*, we summed it over unordered species pairs.

### Cell-level loss

We design a cell-level loss that distinguishes cell types within each species. For species *i*, let 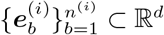 denote the cell embeddings, corresponding to the rows of the cell embedding matrix 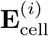, and let 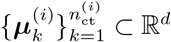 denote the cell-type embeddings, corresponding to the rows of the cell-type embedding matrix 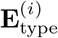. Each cell 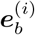 is annotated with a cell-type label 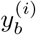. The cell-level loss is computed within each species, pulling each cell embedding towards its corresponding cell-type embedding while pushing it away from all other cell-type embeddings of the same species in the latent space:

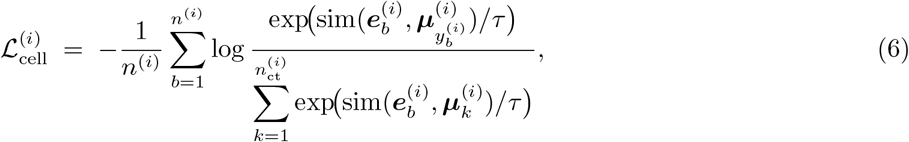

where sim(·, ·) denotes cosine similarity and *τ* is a temperature hyperparameter.

### Objective function and two-stage training

We optimize a weighted sum of the gene- and cell-level losses:

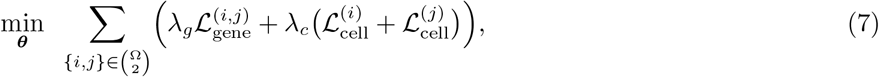

where 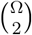 denotes the set of unordered species pairs, ***θ*** denotes all learnable model parameters, and *λ*_*g*_ ≥ 0 and *λ*_*c*_ ≥ 0 balance the two objectives. We adopt a two-stage training strategy. In the first stage, we optimize the gene-level objective only by setting *λ*_*c*_ = 0 to initialize a shared gene embedding space across species. In the second stage, we optimize the full objective with *λ*_*g*_ *>* 0 and *λ*_*c*_ *>* 0.

### Cross-species cell-type tree construction

To obtain a stable cross-species cell-type tree, we aggregated cell-type embeddings from the last *M* training epochs rather than relying on a single checkpoint (in our experiments *M* = 50). For each epoch, we computed pairwise distances between all cell types across species using a cosine-derived distance and averaged the resulting distance matrices across epochs. We then constructed the main tree using Ward’s hierarchical clustering, as implemented in SciPy [46]. To assess the robustness of each internal split, we compared the main tree with the epoch-wise trees and assigned each cluster a fuzzy support score based on its consistency across epochs (Supplementary Note 5).

### Datasets and preprocessing

#### Embryogenesis datasets (frog and zebrafish)

The frog and zebrafish embryogenesis single-cell RNA-seq datasets were originally generated by Briggs et al. [21]. In this study, we used the processed expression matrices and the one-to-one orthologs provided by Zhong et al. [47]. These matrices had already been normalized and log-transformed. We then selected 2,000 highly variable genes separately for each species. The embryogenesis datasets included 63,371 zebrafish cells and 96,935 frog cells, spanning 42 cell types in zebrafish and 36 cell types in frog.

#### Primary motor cortex datasets (human, mouse and marmoset)

The human, mouse and marmoset primary motor cortex single-cell RNA-seq datasets used in this study were downloaded from Biharie et al. [15], based on the original data generated by Bakken et al. [22]. For each species pair, one-to-one orthologs were retrieved from Ensembl BioMart [48] using Ensembl Genes 115 [49]. Starting from raw UMI count matrices, we removed cells with fewer than 200 total UMI counts and genes detected in fewer than 10 cells. Counts were then normalized per cell and log-transformed in Scanpy [50]. We next selected 2,000 highly variable genes separately for each species. After preprocessing, the primary motor cortex datasets comprised 76,533 human cells, 69,279 marmoset cells and 159,738 mouse cells. They were annotated at both low and high resolution, with 20, 22 and 23 low-resolution cell types in human, marmoset and mouse, respectively, and 46 high-resolution cell types in each species, one of which was labeled “NA”.

### Benchmarks

#### Integration methods

We compared CHORD with four baseline data integration methods: SATURN [16], scGen [25], scVI [23] and scANVI [24]. For all methods, we followed the input requirements of the original publications and used default parameters unless otherwise noted. Methods supporting cross-species supervision (scANVI and scGen) used harmonized annotations (Supplementary Note 1).

#### Metrics of integration performance

We evaluated integration performance from two aspects: batch correction and biological conservation. The batch correction score was defined as the average of batch average silhouette width (bASW) and graph connectivity (GC), with species treated as the batch variable. The biological conservation score was defined as the average of four metrics: cell-type average silhouette width (cASW), cell-type normalized mutual information (NMI), cell-type adjusted Rand index (ARI) and trajectory conservation score (Traj). Trajectory conservation was computed only for the frog and zebrafish embryogenesis datasets, which capture developmental processes with a clear temporal ordering of cells (Supplementary Note 3).

## 5 Data availability

The frog and zebrafish embryogenesis single-cell RNA-seq datasets and one-to-one orthologs used in this study are available at https://figshare.com/s/6187811b6c3fae02a4d3. The primary motor cortex single-cell RNA-seq datasets from human, mouse and marmoset used in this study are available at https://zenodo.org/records/11191718. The one-to-one orthologs for human, mouse and marmoset are available at http://www.ensembl.org/biomart.

## 6 Code availability

The source code of CHORD is available on GitHub at https://github.com/zjupgx/CHORD.

## 8 Acknowledgments

We thank the Information Technology Center and State Key Lab of CAD&CG, the Innovation Institute for Artificial Intelligence in Medicine, Zhejiang University, and Shanghai Innovation Institute for the support of computing resources.

