## Supplementary Information for "CHORD: a framework for cross-species single-cell integration across gene, cell and cell-type levels"

#### Supplementary Note 1 Baseline methods

We compared CHORD with four baseline data integration methods: SATURN [1], scGen [2], scVI [3] and scANVI [4]. With respect to input data, scVI, scANVI and SATURN were run on raw counts, whereas scGen was run on the same preprocessed expression matrices used as input to CHORD. For methods that require a shared gene feature space across species (scVI, scANVI and scGen), we first restricted the gene set to one-to-one orthologs and then selected 2000 highly variable genes, treating species as the batch variable. SATURN requires protein-level embeddings as input features. For human, mouse and marmoset, we downloaded protein FASTA sequences from Ensembl (Ensembl Genes 115) and computed protein embeddings using the ESM2 model [5] (`esm2_t33_650M_UR50D`), following the procedure described in the article [1]. For frog and zebrafish, we used the precomputed protein embeddings provided by the SATURN authors [1].

#### Supplementary Note 2 Hyperparameters

We selected the top 2,000 highly variable genes per species during preprocessing. In CHORD, we set the dimension of both gene embeddings and cell-type prototypes to 128, and the hidden dimension of the MLPs was also 128. We included a mutual k-nearest neighbors regularization term with $k = 2$ and set the masking probability to $p = 0.1$. All models were trained with a batch size of 256 and contrastive temperature $\tau = 0.1$ using the Adam optimizer. Training was organized into two stages: in the first stage, we trained for 100 epochs with a learning rate of $1 \times 10^{-3}$ and loss weights $\lambda_g = 1$, $\lambda_c = 0$; in the second stage, we trained for an additional 200 epochs with a learning rate of $5 \times 10^{-3}$ and loss weights $\lambda_g = 10$, $\lambda_c = 0.1$. For low-to-high experiments, we fine-tuned the model for 100 epochs with a learning rate of $5 \times 10^{-3}$ and loss weights $\lambda_g = 0$, $\lambda_c = 0.1$, while freezing the gene embeddings.

#### Supplementary Note 3 Metrics of integration performance

All metrics were computed on the output embeddings of the methods and described as follows:

- **Batch ASW (bASW)**: Quantifies batch mixing when species is treated as the batch label. It was computed following the batch silhouette-based score used in scIB [6] and rescaled to the $[0, 1]$ range, with higher values indicating better integration.
- **Graph connectivity (GC)**: Evaluates how well cells from the same biological population form a single connected neighborhood across species.

- **Cell-type ASW (cASW)**: Quantifies how compact and well-separated annotated cell types are in the integrated embedding, rescaled to the $[0, 1]$ range, with higher values indicating better preservation of cell-type structure.
- **Cell-type NMI (cNMI)**: Measures the agreement between clusters in the integrated space and ground-truth cell-type labels across species. Referring to the implementation of scIB, we computed NMI by applying Leiden clustering with resolution parameters ranging from 0.1 to 2.0 in increments of 0.1, and reported the maximum NMI value over all tested resolutions.
- **Cell-type ARI (cARI)**: Assesses the similarity between clustering and true cell-type labels while correcting for chance agreement. The calculation procedure was the same as for NMI.
- **Trajectory conservation score (Traj)**: Evaluates how well pseudotime structure is preserved after integration. It was computed only for the frog and zebrafish embryogenesis datasets, because these data describe developmental processes with a clear temporal ordering of cells. By contrast, the primary motor cortex datasets do not contain comparable time-resolved trajectories.

#### Supplementary Note 4 Cell-type annotation with unknown cell-type detection

##### Splits of datasets

We utilized the frog and zebrafish embryogenesis datasets to evaluate the performance of cell-type annotation and unknown cell-type detection, because these datasets contain several species-specific cell types. We defined unknown cell-type splits separately for each species. In the frog dataset, the following five cell types were treated as unknown: blastula, myeloid progenitors, pronephric mesenchyme, small secretory cells and Spemann organizer. In the zebrafish dataset, the unknown cell types included dorsal organizer, macrophages, pharyngeal pouch, pluripotent cells and secretory epidermal cells. These cell types were chosen because they are species-specific with sufficient cell numbers. Cell types that were shared across species, together with a subset of species-specific types, were treated as known classes. For each known cell type, we randomly assigned 80% of cells to the training set and the remaining 20% to the test set. Cells belonging to unknown cell types were never used for training and were included only in the test set. Importantly, HVG selection and the calculation of the cross-species gene association matrix in CHORD were performed using only the training data, excluding all test cells, in order to avoid information leakage.

##### Cell-type annotation and unknown cell-type detection

Given a query species $Q$ and a reference species $R$, we treat cells from $Q$ as queries. For CHORD, the references are the cell-type embeddings learned for species $R$, denoted by $\{\boldsymbol{\mu}_k^{(R)}\}_{k=1}^{n_{\text{ct}}^{(R)}}$. For other integration methods, we first obtain integrated embeddings for cells and then train a multinomial logistic regression classifier on embeddings from species $R$ using their cell-type labels; the classifier outputs class probabilities for cells from $Q$. When $Q$ is equal to $R$, this corresponds to within-species classification; when $Q$ is not equal to $R$, this corresponds to cross-species label transfer from species $R$ to species $Q$.

For CHORD, which represents each cell type by a learnable cell-type embedding, we classify cells from a query species $Q$ against a reference species $R$ as follows. Let $\mathbf{E}_{\text{cell}}^{(Q)}$ denote the matrix of cell embeddings for species $Q$, whose $b$-th row is the embedding $\boldsymbol{e}_b^{(Q)}$ of the $b$-th cell. Similarly, let $\mathbf{E}_{\text{type}}^{(R)}$ denote the matrix of cell-type embeddings for species $R$, whose $k$-th row is the cell-type embedding $\boldsymbol{\mu}_k^{(R)}$. We compute cosine similarities, scale them by a temperature parameter $\tau$ ($\tau = 0.1$), and convert them to class probabilities via a softmax:

$$\pi_{bk}^{(Q\to R)} = \frac{\exp\left(\frac{\mathrm{sim}\left(\boldsymbol{e}_b^{(Q)}, \boldsymbol{\mu}_k^{(R)}\right)}{\tau}\right)}{\sum_{j=1}^{n_{\mathrm{ct}}^{(R)}} \exp\left(\frac{\mathrm{sim}\left(\boldsymbol{e}_b^{(Q)}, \boldsymbol{\mu}_j^{(R)}\right)}{\tau}\right)}. \tag{1}$$

Let $\mathcal{K}^{(R)}$ denote the set of known cell types in the reference species. We define the margin (confidence score) for cell $b$ as the maximum predicted class probability over all known cell types:

$$m_b^{(Q\to R)} = \max_{k\in\mathcal{K}^{(R)}} \pi_{bk}^{(Q\to R)}. \tag{2}$$

For the other integration methods, given embeddings for species $Q$ and a classifier trained on embeddings from species $R$, we obtain predicted probabilities $\{\pi_{bk}^{(Q\to R)}\}_{k\in\mathcal{K}^{(R)}}$ for cells from $Q$ in the same way and define the margin $m_b^{(Q\to R)}$ identically as the maximum over known classes.

Unknown cell-type detection is implemented by sweeping 200 evenly spaced rejection thresholds $\theta \in [0, 1]$ over the margin. For a given $\theta$, the predicted label for cell $b$ is:

$$\hat{y}_b^{(Q\to R)}(\theta) = \begin{cases} \arg\max\limits_{k\in\mathcal{K}^{(R)}} \pi_{bk}^{(Q\to R)}, & \text{if } m_b^{(Q\to R)} \geq \theta, \\ \text{Unknown}, & \text{if } m_b^{(Q\to R)} < \theta. \end{cases} \tag{3}$$

Let $y_b$ be the true cell-type label, and let $\mathcal{I}_{\mathrm{known}}$ and $\mathcal{I}_{\mathrm{unknown}}$ denote the index sets of cells whose true labels belong to the known and unknown classes, respectively. The correct classification rate on known cells ($\mathrm{CCR}_{\mathrm{known}}$) at threshold $\theta$ is defined as:

$$\mathrm{CCR}_{\mathrm{known}}(\theta) = \frac{1}{|\mathcal{I}_{\mathrm{known}}|} \sum_{b\in\mathcal{I}_{\mathrm{known}}} \mathbf{1}\left[\, \hat{y}_b^{(Q\to R)}(\theta) = y_b \,\right], \tag{4}$$

which is the proportion of known cells that are correctly classified and not rejected. The recall on unknown cells at threshold $\theta$ is defined as:

$$\mathrm{Recall}_{\mathrm{unknown}}(\theta) = \frac{1}{|\mathcal{I}_{\mathrm{unknown}}|} \sum_{b\in\mathcal{I}_{\mathrm{unknown}}} \mathbf{1}\left[\, \hat{y}_b^{(Q\to R)}(\theta) = \text{Unknown} \,\right], \tag{5}$$

which is the proportion of truly unknown cells that are correctly rejected as unknown. By varying the threshold $\theta$, we obtain a curve that characterizes the trade-off between maintaining accuracy on known cell types and correctly rejecting unknown cell types.

##### Clustering quality of withheld cell types

To evaluate the clustering quality of withheld cell types, we restricted the analysis to cells whose true labels belonged to the predefined set of withheld cell types. We used ARI and NMI to quantify agreement between clustering results and the ground-truth withheld cell-type labels. We applied Leiden clustering on the integrated embedding over resolutions from 0.1 to 2.0 in increments of 0.1; for each resolution, we computed ARI and NMI between the predicted clusters and the ground-truth labels. We then selected the resolution that maximized

ARI and reported both ARI and NMI at this resolution. To assess intrinsic cluster quality, we computed the Davies–Bouldin index and ASW using the ground-truth labels as cluster assignments.

#### Supplementary Note 5 Cross-species cell-type tree construction

To construct a stable cell-type tree, we do not rely on a single checkpoint. Instead, we collect the cell-type embeddings from the last $M$ training epochs (in our experiments $M = 50$). We index cell types by a pair $(i, s)$, where $i$ denotes the species and $s$ denotes a cell type within species $i$. To distinguish epoch snapshots, we write $\boldsymbol{\mu}_s^{(i)}[r]$ for the embedding of cell type $s$ in species $i$ taken from the $r$-th epoch snapshot, with $r = 1, \ldots, M$. For two cell types $(i, s)$ and $(j, t)$ at snapshot $r$, we define their distance as:

$$d^{(r)}\big((i,s),(j,t)\big) = \sqrt{2\left(1 - \cos\left(\boldsymbol{\mu}_s^{(i)}[r], \boldsymbol{\mu}_t^{(j)}[r]\right)\right)}. \tag{6}$$

For each snapshot $r$, these distances define an $N \times N$ pairwise distance matrix:

$$D^{(r)}_{(i,s),(j,t)} = d^{(r)}\big((i,s),(j,t)\big), \tag{7}$$

where $N$ is the total number of cell types across species. To construct the cell-type tree, we average these matrices:

$$\bar{D} = \frac{1}{M}\sum_{r=1}^{M} D^{(r)}, \tag{8}$$

and apply hierarchical clustering with Ward linkage.

To quantify how consistently each split appears across the $M$ epoch-wise trees, we assign a fuzzy support value in $[0, 1]$ to every internal cluster in the main cell-type tree. For a cluster $C$ in the main tree and each epoch snapshot $r$, we identify the most similar cluster in the $r$-th epoch-wise tree using an F1 similarity over the corresponding sets of cell-type labels, and denote the resulting similarity by $s^{(r)}(C)$. Specifically, for two clusters $A$ and $B$, we define:

$$\mathrm{F1}(A, B) = \frac{2|A \cap B|}{|A| + |B|}. \tag{9}$$

We then average these similarities across epoch snapshots, ignoring values below a fixed similarity threshold $\kappa$:

$$\mathrm{support}(C) = \frac{1}{M}\sum_{r=1}^{M} \mathbf{1}\big(s^{(r)}(C) \geq \kappa\big)\, s^{(r)}(C), \tag{10}$$

where $\mathbf{1}(\cdot)$ denotes the indicator function. We set $\kappa = 0.8$ in our experiments. We interpret values $< 0.5$ as low, 0.5–0.8 as intermediate, and $\geq 0.8$ as high support.

#### Supplementary Note 6 Tracing gene expression dynamics on the cell-type tree

To trace the branch-specific dynamics of a given one-to-one ortholog $g$, we mapped its expression profile onto the inferred cell-type tree. In this binary tree, leaves represent species-specific cell types, and internal nodes represent progressively broader hierarchical groupings of related cell types. For each species $i$ and cell type $s$, let $h_{g,s}^{(i)}$ denote the average log-transformed normalized expression of gene $g$ across all cells annotated as $s$ in species $i$. We assigned this value to the corresponding leaf $\ell$ of the tree representing the species-specific cell type $(i, s)$:

$$e_\ell = h_{g,s}^{(i)}. \tag{11}$$

To infer internal-node expression summaries, we aggregated expression levels across all descendant leaves. For an internal node $u$, let $\mathcal{L}(u)$ be the set of all leaves in the subtree rooted at $u$. We defined the node-level expression $e_u$ as the mean expression of its descendants:

$$e_u = \frac{1}{|\mathcal{L}(u)|} \sum_{\ell \in \mathcal{L}(u)} e_\ell. \tag{12}$$

Finally, to quantify branch-specific expression changes, we computed the difference in expression between every non-root node $v$ and its parent node $p(v)$:

$$\Delta e_v = e_v - e_{p(v)}. \tag{13}$$

A positive $\Delta e_v$ indicates that the ortholog is upregulated in node $v$ relative to its parent (gain), whereas a negative value indicates downregulation (loss). These $\Delta e_v$ values were used to color the branches of the tree, providing a branch-resolved view of gene expression dynamics along the cell-type tree.

#### Supplementary Note 7 Hierarchical fine-tuning

We compared three training settings that differ in the annotation resolution used during training: low-resolution-only, high-resolution-only, and low-to-high. Because the low-resolution-only setting does not directly produce the high-resolution representations required for these evaluations, we derived high-resolution cell-type embeddings from the low-resolution-only model as follows. Starting from the model trained with low-resolution labels, we initialized a prototype for each high-resolution cell type, kept the gene embeddings and MLPs fixed, and optimized only these prototypes with the cell-level loss. This procedure yields high-resolution cell-type embeddings for the low-resolution-only model, enabling a fair comparison in the downstream tests.

To evaluate biological plausibility, we performed two tests: consistency with the low-resolution structure and cross-species transfer of high-resolution labels. For the consistency analysis, we constructed a mapping from high-resolution cell-type labels to low-resolution evaluation groups based on the core cell-type name in the high-resolution annotations, excluding types whose label was "NA" (Supplementary Table 1). We then cut the high-resolution cell-type tree to obtain $K$ clusters, with $K = 20$ (the number of low-resolution evaluation groups). The resulting clusters define low-resolution labels induced from the high-resolution tree. We quantified this consistency by computing NMI and ARI between these induced low-resolution labels and the low-resolution evaluation groups. For the cross-species transfer of high-resolution labels, we applied a prototype-based label transfer strategy similar to that described in Supplementary Note 3. Query cells from species $Q$ were compared against the high-resolution cell-type embeddings of the reference species $R$, and each cell was assigned to the most similar high-resolution cell-type embedding in $R$. We evaluated cross-species annotation performance at high resolution using accuracy and macro-averaged F1 score.

### Supplementary Note 8 Developmental alignment

##### Developmental time alignment between zebrafish and frog

Because the zebrafish and frog datasets use different time labels, we first established a shared developmental time based on the CHORD cell embeddings. We constructed a similarity matrix using symmetric 1-nearest-neighbor cosine distances between frog and zebrafish time points. Let $\mathcal{F}_{t_f}$ denote the set of frog cells at frog time $t_f$ with embeddings $\{\boldsymbol{e}_b^{(f)} : b \in \mathcal{F}_{t_f}\}$, and let $\mathcal{Z}_{t_z}$ denote the set of zebrafish cells at zebrafish time $t_z$ with embeddings $\{\boldsymbol{e}_c^{(z)} : c \in \mathcal{Z}_{t_z}\}$. For each pair $(t_f, t_z)$, we first computed directional 1-nearest-neighbor cosine distances:

$$d_{f \to z}(t_f, t_z) = \frac{1}{|\mathcal{F}_{t_f}|} \sum_{b \in \mathcal{F}_{t_f}} \min_{c \in \mathcal{Z}_{t_z}} \Big[1 - \cos\big(\boldsymbol{e}_b^{(f)},\, \boldsymbol{e}_c^{(z)}\big)\Big] \tag{14}$$

$$d_{z \to f}(t_f, t_z) = \frac{1}{|\mathcal{Z}_{t_z}|} \sum_{c \in \mathcal{Z}_{t_z}} \min_{b \in \mathcal{F}_{t_f}} \Big[1 - \cos\big(\boldsymbol{e}_c^{(z)},\, \boldsymbol{e}_b^{(f)}\big)\Big]. \tag{15}$$

We then defined a symmetric distance as the average of the two directions:

$$d_{\text{sym}}(t_f, t_z) = \frac{1}{2}\Big(d_{f \to z}(t_f, t_z) + d_{z \to f}(t_f, t_z)\Big), \tag{16}$$

and converted it to a similarity score:

$$S(t_f, t_z) = 1 - d_{\text{sym}}(t_f, t_z). \tag{17}$$

The similarity matrix $\mathbf{S}$ has entries:

$$\mathbf{S}_{f,z} = S(t_f, t_z), \tag{18}$$

with frog times $t_f$ as rows and zebrafish times $t_z$ as columns. In the resulting similarity matrix, rows correspond to frog developmental stages and columns to zebrafish time points. For the initial alignment, we applied a row-wise argmax, assigning to each frog stage the zebrafish time point with the highest similarity. To obtain a smooth, strictly increasing alignment curve, we linearly interpolated between the assignments for Stage 10 and Stage 12 in zebrafish time and assigned Stage 11 an intermediate value (7 hpf).

##### Per-cell aligned transcriptional tempo

Per-cell transcriptional tempo was computed in the CHORD cell embedding as forward movement per unit time along the aligned developmental time. Let $\mathcal{F}_t$ denote the set of frog cells whose aligned time is $t$ (in zebrafish hours post-fertilization) with embeddings $\{\boldsymbol{e}_b^{(f)} : b \in \mathcal{F}_t\}$, and let $\mathcal{Z}_t$ denote the set of zebrafish cells at aligned time $t$ with embeddings $\{\boldsymbol{e}_c^{(z)} : c \in \mathcal{Z}_t\}$.

For each cell, we used the nearest available next aligned time point $t'$ within the same species to define its transcriptional tempo. For example, for a frog cell $\boldsymbol{e}_b^{(f)} \in \mathcal{F}_t$ with next aligned time point $t' > t$, we defined its transcriptional tempo as:

$$r_b^{(f)}(t \to t') \;=\; \frac{1}{\Delta t} \min_{b' \in \mathcal{F}_{t'}} \Big[1 - \cos\big(\boldsymbol{e}_b^{(f)}, \boldsymbol{e}_{b'}^{(f)}\big)\Big], \tag{19}$$

where $\Delta t = t' - t$ is the time interval between the two aligned time points. An analogous definition was used for zebrafish cells.

### Supplementary Note 9 Gene analysis

#### Orthology-based evaluation of gene embeddings

To assess whether the gene embeddings recover orthology relationships, we compared the similarity of one-to-one ortholog pairs to randomly paired genes across species. For each species pair, we extracted gene embeddings of all annotated one-to-one orthologs and computed their cosine similarities. As a baseline, we sampled an equal number of random cross-species gene pairs from the two species while excluding true one-to-one ortholog pairs, and computed cosine similarities for these random pairs in the same way. We then contrasted the two similarity distributions using a one-sided Wilcoxon rank-sum test (alternative hypothesis: ortholog pairs have higher similarity than random pairs).

#### Gene-embedding-informed analysis of protein–protein interaction networks

We used protein–protein interaction (PPI) networks to assess whether the learned gene embeddings recover known functional neighborhoods within each species and whether these relationships are consistent between human and mouse. For each gene $g$, we defined an embedding-based neighborhood $\mathcal{N}^{\text{emb}}(g)$ as the set of genes with the highest cosine similarities to $g$, excluding $g$ itself. PPI neighborhoods $\mathcal{N}^{\text{PPI}}(g)$ were obtained from the STRING database [7] for the corresponding species, after removing self-loops. For each species, we defined the background gene universe $\mathcal{B}$ as the set of genes present in both the embedding space and the species-specific PPI network. Within $\mathcal{B}$, we quantified the agreement between $\mathcal{N}^{\text{emb}}(g)$ and $\mathcal{N}^{\text{PPI}}(g)$ for each gene $g$. Let $M = |\mathcal{B}|$ denote the number of background genes, $n = |\mathcal{N}^{\text{PPI}}(g) \cap \mathcal{B}|$ the number of PPI partners of $g$ in the background, $N = |\mathcal{N}^{\text{emb}}(g) \cap \mathcal{B}|$ the number of embedding neighbours in the background, and $k = |\mathcal{N}^{\text{PPI}}(g) \cap \mathcal{N}^{\text{emb}}(g) \cap \mathcal{B}|$ the size of their overlap. We modeled:

$$X \sim \text{Hypergeom}(M, n, N), \tag{20}$$

and computed a one-sided $P$ value:

$$P_{\text{overlap}} = P(X \geq k), \tag{21}$$

assigning $P_{\text{overlap}} = 1$ when $n = 0$, $N = 0$, or $k = 0$. Benjamini–Hochberg correction was applied per species to obtain false discovery rate (FDR)-adjusted $q$ values.

To prioritize associations jointly supported in human and mouse, we joined the human and mouse tables via the one-to-one ortholog mapping, yielding for each ortholog pair $(g_{\text{human}}, g_{\text{mouse}})$ a pair of per-species overlap $P$ values $(p_{\text{human}}, p_{\text{mouse}})$ whenever both were available. We performed a cross-species meta-analysis using Fisher's method. For each pair, we computed:

$$X = -2 \sum_{i \in \{\text{human},\text{mouse}\}} \ln p_i, \tag{22}$$

which follows a $\chi^2$ distribution with 4 degrees of freedom under the null hypothesis of no enrichment in either species. The resulting meta-analysis $P$ values were adjusted for multiple testing across all tested ortholog pairs using the Benjamini–Hochberg procedure, yielding $q_{\text{meta}}$.

##### Gradient-based gene importance scores

For each species $i$ and a given target cell type, we quantified gene-level importance using a gradient–input attribution scheme, in which the importance score reflects the marginal contribution of each gene to the predicted probability of that cell type. Let $\mathcal{C}^{(i)}$ denote the set of cells from species $i$ belonging to this target cell type, and let $\boldsymbol{x}_b^{(i)} \in \mathbb{R}^{|\mathcal{G}_{\text{HVG}}^{(i)}|}$ be the input expression vector of cell $b \in \mathcal{C}^{(i)}$ over HVGs $\mathcal{G}_{\text{HVG}}^{(i)}$. We constructed a scalar objective as the mean cosine similarity between the embeddings of the target cells and their cell type:

$$\ell^{(i)} = \frac{1}{|\mathcal{C}^{(i)}|} \sum_{b \in \mathcal{C}^{(i)}} \text{sim}\big(\boldsymbol{e}_b^{(i)}, \boldsymbol{\mu}^{(i)}\big), \tag{23}$$

where $\text{sim}(\cdot, \cdot)$ denotes the cosine similarity, and $\boldsymbol{e}_b^{(i)}$ and $\boldsymbol{\mu}^{(i)}$ are the cell and cell-type embeddings.

We then computed gradients of this objective with respect to the input expression vectors:

$$\boldsymbol{r}_b^{(i)} = \frac{\partial \ell^{(i)}}{\partial \boldsymbol{x}_b^{(i)}} \in \mathbb{R}^{|\mathcal{G}_{\text{HVG}}^{(i)}|}, \quad b \in \mathcal{C}^{(i)}, \tag{24}$$

and used a gradient–input product to define per-gene importance magnitudes. For gene $g \in \{1, \ldots, |\mathcal{G}_{\text{HVG}}^{(i)}|\}$, the importance in species $i$ was given by:

$$m_g^{(i)} = \sum_{b \in \mathcal{C}^{(i)}} \big| r_{b,g}^{(i)} \, x_{b,g}^{(i)} \big|, \tag{25}$$

where $r_{b,g}^{(i)}$ and $x_{b,g}^{(i)}$ are the $g$-th entries of $\boldsymbol{r}_b^{(i)}$ and $\boldsymbol{x}_b^{(i)}$, respectively. To facilitate comparison of importance values across genes within each species, we applied a z-score normalization, providing a rank-like measure of relative gene importance $z_g^{(i)}$ within each species.

#### Supplementary Note 10 Stochastic gene masking in CHORD

CHORD introduces a stochastic gene expression mask $\mathbf{M}^{(i)} \in \{0,1\}^{n^{(i)} \times |\mathcal{G}_{\text{HVG}}^{(i)}|}$, where $n^{(i)}$ is the number of cells for species $i$. The stochastic mask acts as feature dropout and encourages robust representations. Each element $\mathbf{M}_{ng}^{(i)}$ of the mask is independently sampled from a Bernoulli distribution to indicate whether a gene expression is kept or masked:

$$\mathbf{M}_{ng}^{(i)} \sim \text{Bernoulli}(1 - p), \tag{26}$$

where $p$ is the probability of masking a gene. The mask is then applied to the preprocessed single-cell gene expression matrix, denoted $\mathbf{X}'^{(i)} \in \mathbb{R}^{n^{(i)} \times |\mathcal{G}_{\text{HVG}}^{(i)}|}$, via an element-wise (Hadamard) product:

$$\tilde{\mathbf{X}}^{(i)} = \mathbf{X}'^{(i)} \odot \mathbf{M}^{(i)}. \tag{27}$$

#### Supplementary Note 11 Construction of the cross-species cell-type correspondence matrix in CHORD

For a species pair $(i, j)$, we denote the set of one-to-one ortholog pairs by:

$$\mathcal{O}_{1:1}^{(i,j)} = \{\, (g_i, g_j) \mid g_i \text{ and } g_j \text{ form a one-to-one ortholog} \,\}. \tag{28}$$

First, we calculate a cell-type-averaged expression matrix $\mathbf{H}^{(i)}$ over $\mathcal{G}_{\mathrm{HVG}}^{(i)}$:

$$\mathbf{H}^{(i)} = [h_{g,s}^{(i)}] \in \mathbb{R}^{\left|\mathcal{G}_{\mathrm{HVG}}^{(i)}\right| \times n_{\mathrm{ct}}^{(i)}} \tag{29}$$

$$h_{g,s}^{(i)} = \frac{1}{|C_s^{(i)}|} \sum_{x \in C_s^{(i)}} \mathrm{expr}_x(g), \tag{30}$$

where $g \in \mathcal{G}_{\mathrm{HVG}}^{(i)}$ indexes rows of $\mathbf{H}^{(i)}$, and $C_s^{(i)}$ denotes the set of cells from species $i$ annotated as cell type $s$; $\mathrm{expr}_x(g)$ denotes the entry of the preprocessed matrix $\mathbf{X'}^{(i)}$ for cell $x$ and gene $g$. Second, for a species pair $(i, j)$, we define the anchor ortholog pairs:

$$\mathcal{O}_{A}^{(i,j)} \;=\; \{(g_i, g_j) \in \mathcal{O}_{1:1}^{(i,j)} \;\mid\; g_i \in \mathcal{G}_{\mathrm{HVG}}^{(i)} \;\wedge\; g_j \in \mathcal{G}_{\mathrm{HVG}}^{(j)}\}. \tag{31}$$

Let $\pi_i(\mathcal{O}_A^{(i,j)}) = \{\, g_i \mid (g_i, g_j) \in \mathcal{O}_A^{(i,j)} \,\}$ denote the anchor genes in species $i$, and define $\pi_j(\mathcal{O}_A^{(i,j)})$ analogously. For each cell type $s$ in species $i$, we extract the anchor expression vector by restricting $\mathbf{H}^{(i)}$ to genes in $\pi_i(\mathcal{O}_A^{(i,j)})$:

$$\mathbf{a}_s^{(i)} = \big(h_{g,s}^{(i)}\big)_{g \in \pi_i(\mathcal{O}_A^{(i,j)})} \in \mathbb{R}^{|\pi_i(\mathcal{O}_A^{(i,j)})|}. \tag{32}$$

Similarly, $\mathbf{a}_t^{(j)}$ is defined by restricting $\mathbf{H}^{(j)}$ to $\pi_j(\mathcal{O}_A^{(i,j)})$. We compute a similarity matrix $\mathbf{P}^{(i,j)} \in \mathbb{R}^{n_{\mathrm{ct}}^{(i)} \times n_{\mathrm{ct}}^{(j)}}$ with entries:

$$\mathbf{P}_{st}^{(i,j)} = \mathrm{sim}\big(\mathbf{a}_s^{(i)}, \mathbf{a}_t^{(j)}\big), \tag{33}$$

where $\mathrm{sim}(\cdot,\cdot)$ denotes cosine similarity. For each cell type $s$ in species $i$, let $\mathcal{N}_k^{i\to j}(s)$ be the indices of the top-$k$ cell types in species $j$ with the highest similarities in row $s$ of $\mathbf{P}^{(i,j)}$:

$$\mathcal{N}_k^{i\to j}(s) = \arg\mathrm{top}_k\big(\mathbf{P}_{s,\cdot}^{(i,j)}\big). \tag{34}$$

Analogously, for each cell type $t$ in species $j$, let $\mathcal{N}_k^{j\to i}(t)$ be the indices of the top-$k$ cell types in species $i$ with the highest similarities in column $t$ of $\mathbf{P}^{(i,j)}$:

$$\mathcal{N}_k^{j\to i}(t) = \arg\mathrm{top}_k\big(\mathbf{P}_{\cdot,t}^{(i,j)}\big). \tag{35}$$

We define $\mathbf{C}^{(i,j)}$ by retaining only mutual $k$-nearest-neighbor pairs:

$$\mathbf{C}_{st}^{(i,j)} = \begin{cases} \dfrac{1}{k}, & \text{if } t \in \mathcal{N}_k^{i\to j}(s) \text{ and } s \in \mathcal{N}_k^{j\to i}(t) \\[2ex] 0, & \text{otherwise.} \end{cases} \tag{36}$$

#### Supplementary Note 12 Element-wise bound for $\mathbf{S}^{(i,j)}_{\mathbf{cross}}$

We provide a short derivation showing that each entry of $\mathbf{S}^{(i,j)}_{\text{cross}}$ lies in $[-1, 1]$. We first bound the spectral norm of $\mathbf{C}^{(i,j)}$. By construction, each row and each column of $\mathbf{C}^{(i,j)}$ contains at most $k$ nonzero entries, and each nonzero entry is equal to $1/k$. Therefore, every row sum and every column sum is at most 1, which implies:

$$\left\|\mathbf{C}^{(i,j)}\right\|_1 \leq 1, \qquad \left\|\mathbf{C}^{(i,j)}\right\|_\infty \leq 1, \qquad 0 \leq \mathbf{C}^{(i,j)}_{ab} \leq \frac{1}{k} \leq 1. \tag{37}$$

Using the classical inequality relating the spectral norm $\|\cdot\|_2$ to the induced 1- and $\infty$-norms:

$$\|A\|_2 \;\leq\; \sqrt{\|A\|_1\,\|A\|_\infty} \quad \text{for any matrix } A, \tag{38}$$

we obtain:

$$\left\|\mathbf{C}^{(i,j)}\right\|_2 \leq 1\ . \tag{39}$$

Next, recall the definition of $\mathbf{S}^{(i,j)}_{\text{cross}}$:

$$\mathbf{S}^{(i,j)}_{\text{cross}} \;=\; \widehat{\mathbf{H}}^{(i)}\mathbf{C}^{(i,j)}\left(\widehat{\mathbf{H}}^{(j)}\right)^{\mathsf{T}} \;\in \mathbb{R}^{|\mathcal{G}^{(i)}_{\text{HVG}}|\times|\mathcal{G}^{(j)}_{\text{HVG}}|}, \qquad s^{(i,j)}_{ab} \;=\; \left[\mathbf{S}^{(i,j)}_{\text{cross}}\right]_{ab}, \tag{40}$$

where $\widehat{\mathbf{h}}^{(i)}_a$ and $\widehat{\mathbf{h}}^{(j)}_b$ denote the $a$-th and $b$-th rows of $\widehat{\mathbf{H}}^{(i)}$ and $\widehat{\mathbf{H}}^{(j)}$, respectively. By construction, each row vector is $\ell_2$-normalized:

$$\left\|\widehat{\mathbf{h}}^{(i)}_a\right\|_2 \;=\; \left\|\widehat{\mathbf{h}}^{(j)}_b\right\|_2 \;=\; 1. \tag{41}$$

We can then bound each entry $s^{(i,j)}_{ab}$ using the spectral norm of $\mathbf{C}^{(i,j)}$:

$$\left|s^{(i,j)}_{ab}\right| \;=\; \left|\widehat{\mathbf{h}}^{(i)}_a\mathbf{C}^{(i,j)}\left(\widehat{\mathbf{h}}^{(j)}_b\right)^{\mathsf{T}}\right| \;\leq\; \left\|\widehat{\mathbf{h}}^{(i)}_a\right\|_2\left\|\mathbf{C}^{(i,j)}\right\|_2\left\|\widehat{\mathbf{h}}^{(j)}_b\right\|_2 \;\leq\; 1\cdot 1\cdot 1 \;=\; 1. \tag{42}$$

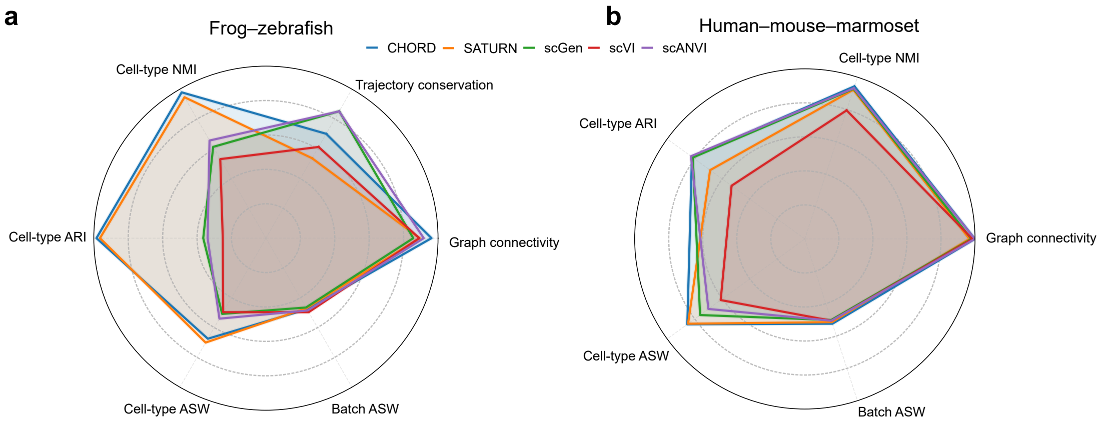


Supplementary Figure 1: Benchmarking cross-species integration methods. Radar plots summarizing performance of CHORD, SATURN, scGen, scVI and scANVI for (a) frog–zebrafish and (b) human–mouse–marmoset datasets. Axes correspond to cell-type NMI, cell-type ARI, cell-type ASW, graph connectivity and batch ASW, with an additional trajectory conservation metric for the frog–zebrafish setting; in all cases, larger values indicate better performance.

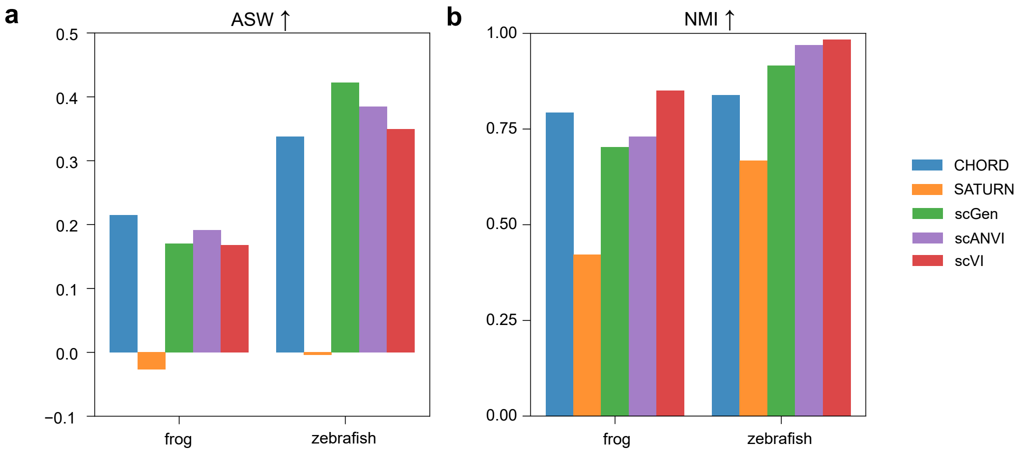


Supplementary Figure 2: Quantification of unknown cell-type detection. We clustered only the unknown cells and evaluated the results with (a) average silhouette width (ASW), which measures cluster compactness and separation (higher is better), and (b) normalized mutual information (NMI) between the inferred clusters and the ground-truth novel cell-type labels (higher is better).

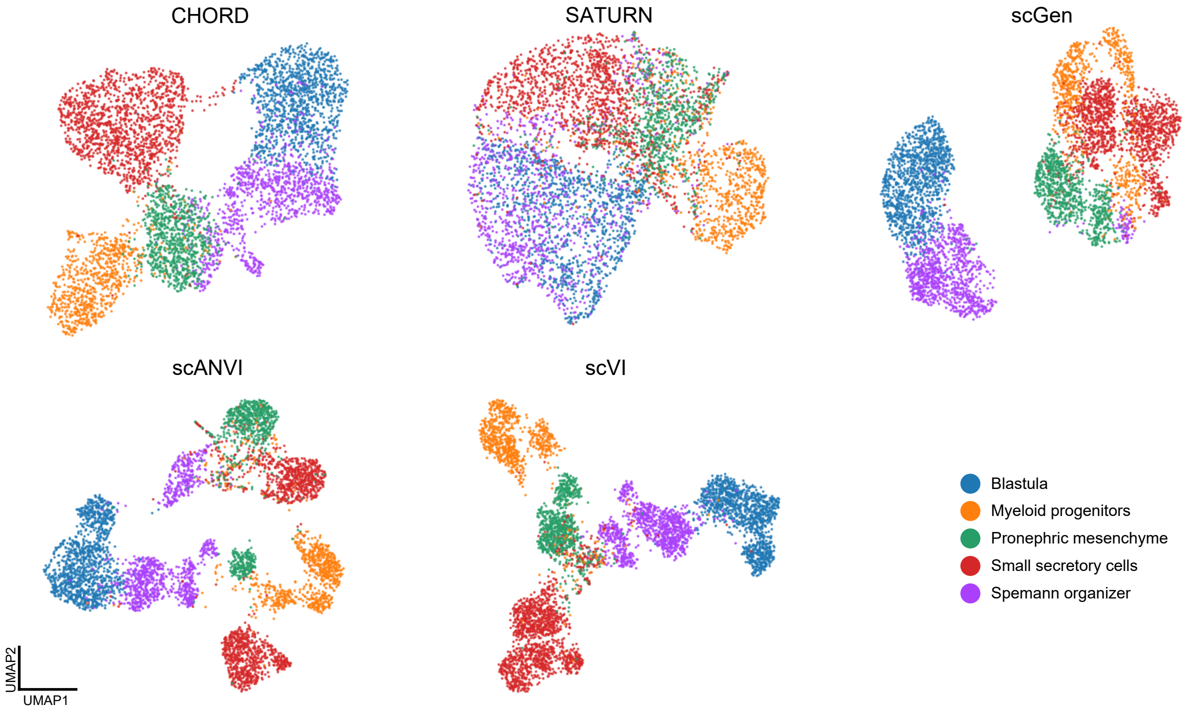


Supplementary Figure 3: UMAP visualizations of unknown frog cell types, where cells withheld as unknown during training are shown for each method (CHORD, SATURN, scGen, scANVI, scVI), colored according to their ground-truth cell-type labels (blastula, myeloid progenitors, pronephric mesenchyme, small secretory cells, Spemann organizer).

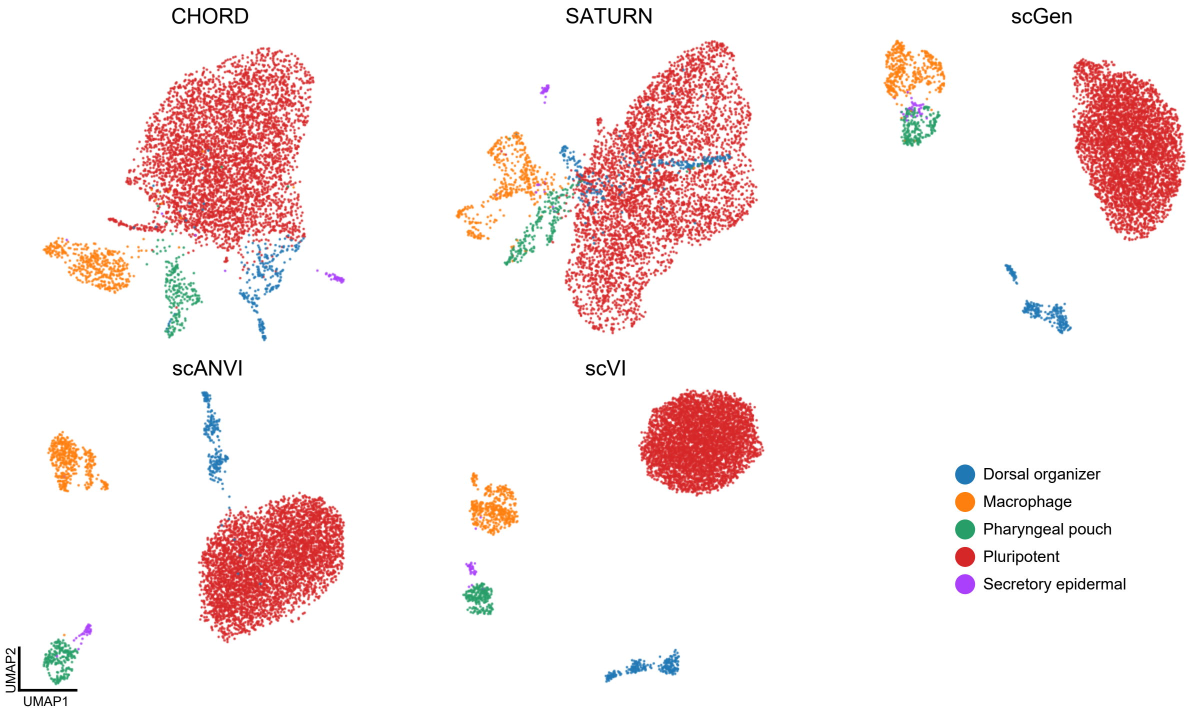


Supplementary Figure 4: UMAP visualizations of unknown zebrafish cell types. Cells withheld as unknown during training are shown for each method (CHORD, SATURN, scGen, scANVI, scVI), colored according to their ground-truth cell-type labels (dorsal organizer, macrophage, pharyngeal pouch, pluripotent, secretory epidermal).

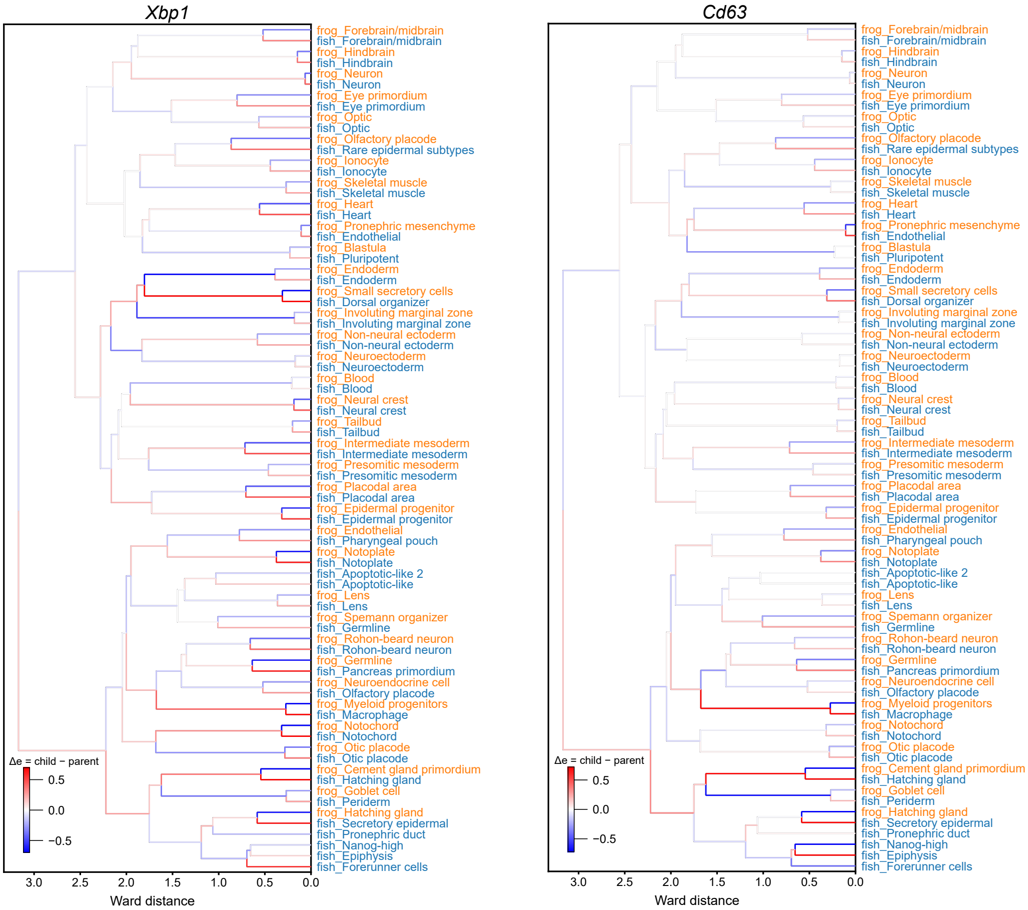


Supplementary Figure 5: Branch-wise changes in *Xbp1* and *Cd63* ortholog expression mapped onto the cell-type tree. For each gene, branch colors indicate the change in mean expression between a parent cluster and each of its children (child minus parent). Red and blue denote increases and decreases in expression.

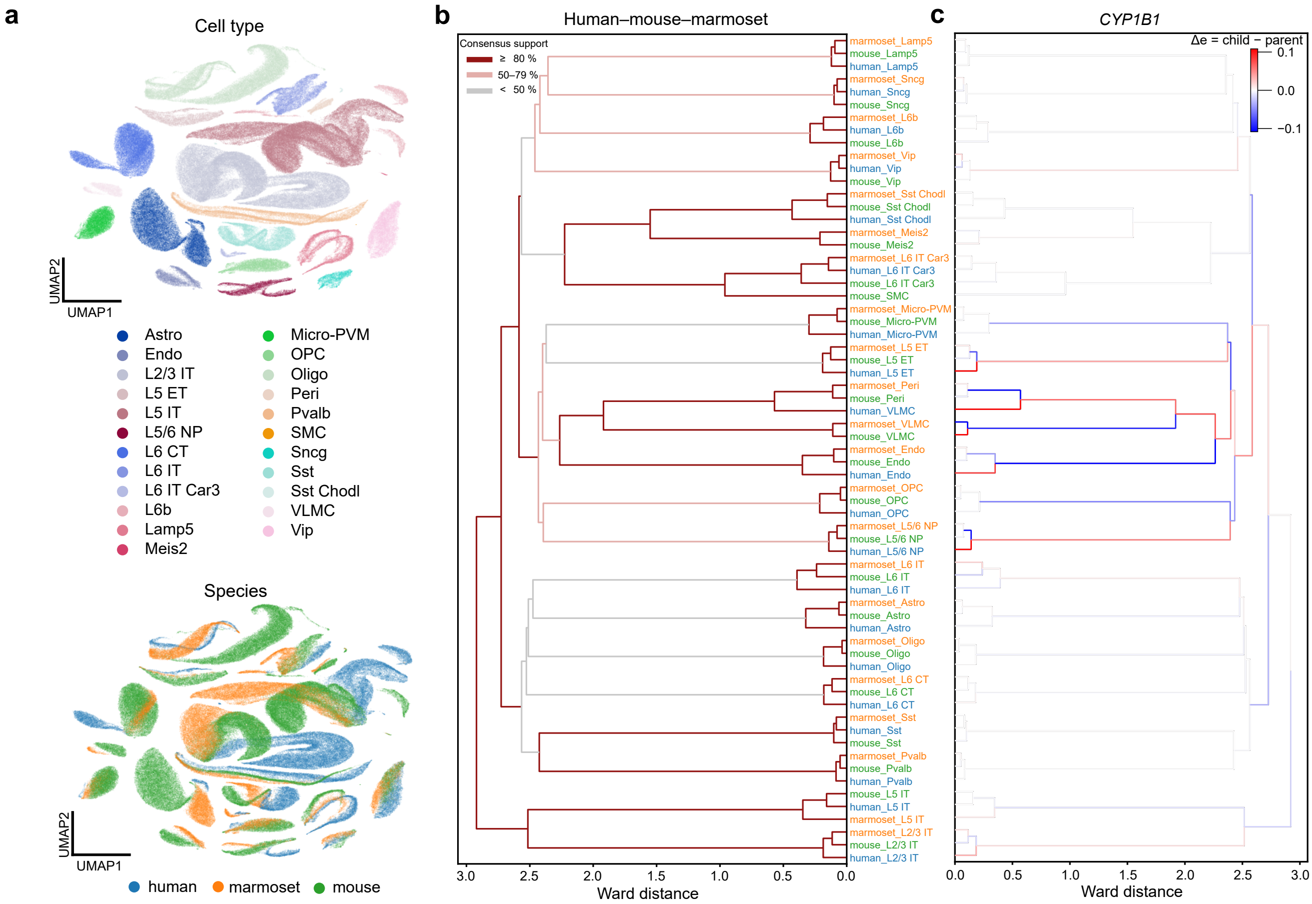


Supplementary Figure 6: CHORD integrates human, mouse and marmoset motor cortex atlases. (a) UMAP visualization of CHORD cell embeddings, colored by cell type (top) and by species (bottom). (b) Cross-species cell-type tree inferred by CHORD. The x-axis shows the Ward distance and leaf labels denote species and cell types. Line colors denote three levels of consensus support. (c) Branch-wise changes in *CYP1B1* ortholog expression mapped onto the cell-type tree in (b) (same topology, mirrored for visualization).

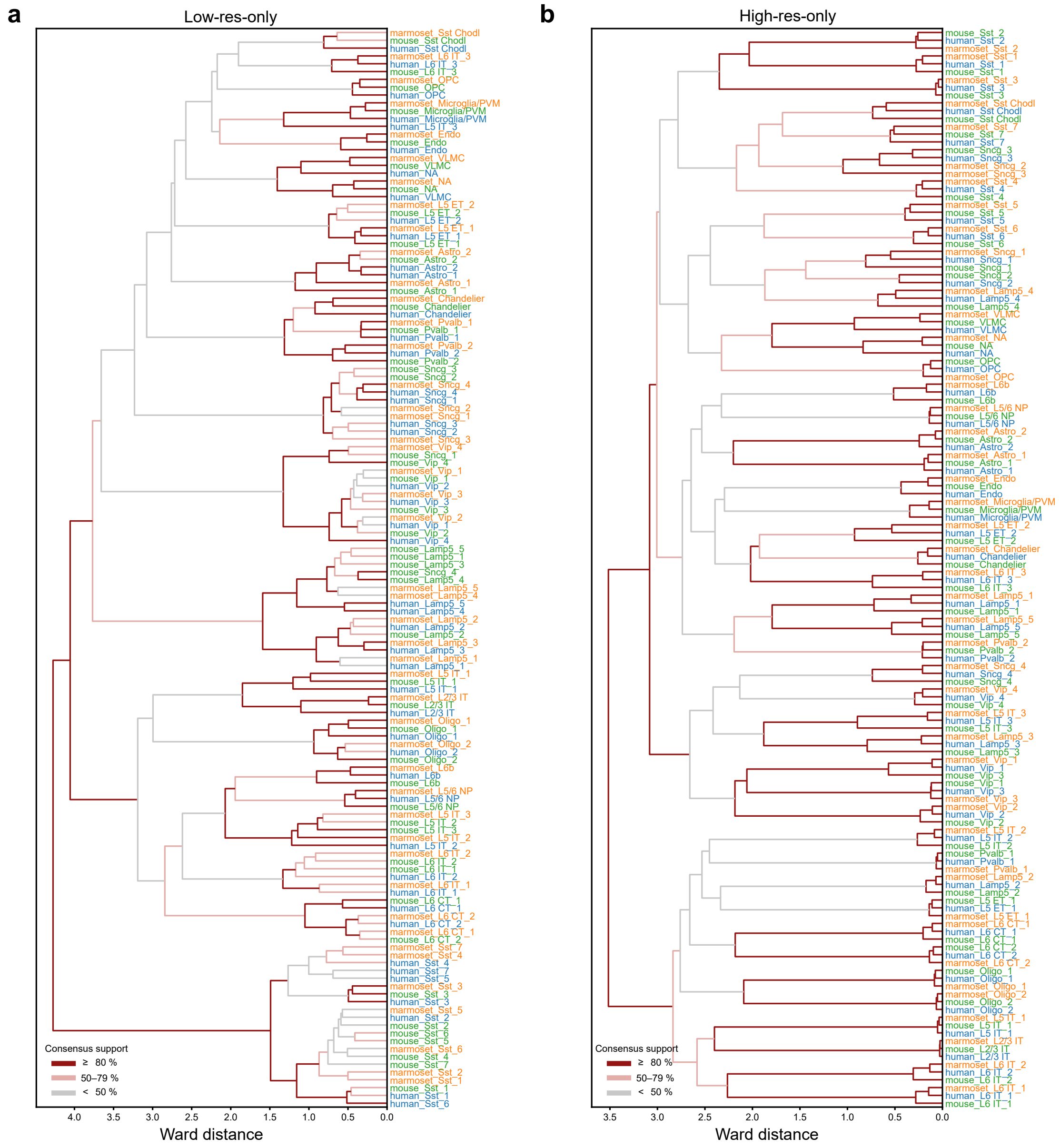


Supplementary Figure 7: Cross-species cell-type trees under different training strategies. The trees inferred (a) from the low-resolution-only model and (b) from the high-resolution-only model.

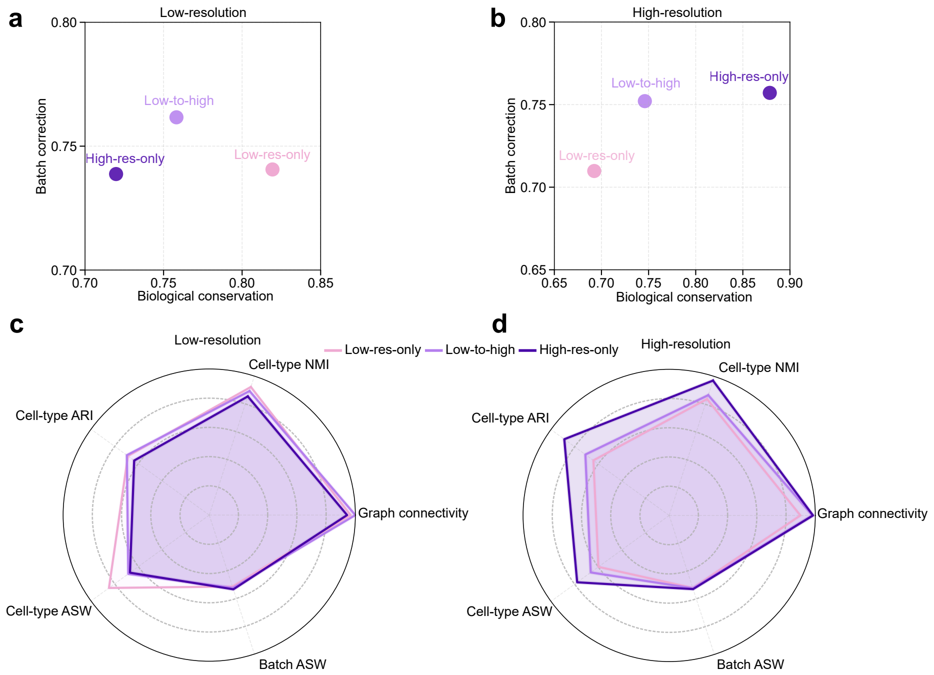


Supplementary Figure 8: Integration performance of different training strategies in CHORD. (a,b) Biological conservation versus batch correction at the (a) low-resolution and (b) high-resolution annotation levels for the low-resolution-only, high-resolution-only and low-to-high models. (c,d) Radar plots summarizing graph connectivity, cell-type NMI, cell-type ARI, cell-type ASW and batch ASW at the (c) low-resolution and (d) high-resolution annotation levels for the three training strategies.

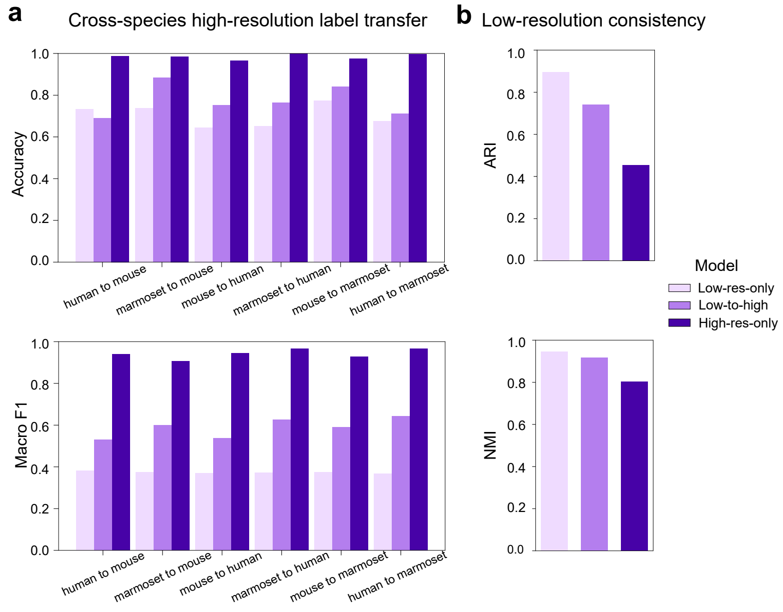


Supplementary Figure 9: Biological evaluation of cross-species cell-type trees across different training strategies. (a) Accuracy and macro F1 for high-resolution cross-species label transfer between all pairs of human, mouse and marmoset. (b) Consistency with the low-resolution structure, quantified by ARI and NMI.

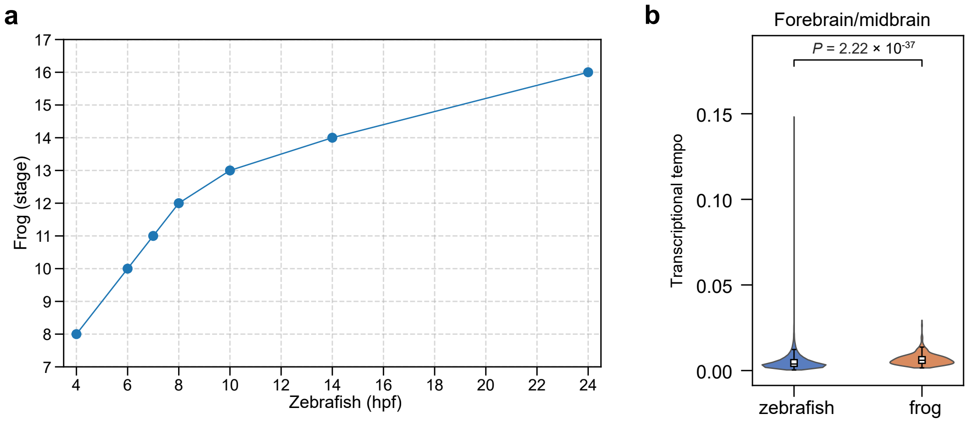


Supplementary Figure 10: Developmental timing differences between zebrafish and frog in the CHORD embedding. (a) Interpolated mapping between zebrafish hours post-fertilization (hpf) and frog stages within the retained window (Stages 8–16). (b) Distributions of transcriptional tempo for zebrafish and frog cells from forebrain/midbrain.

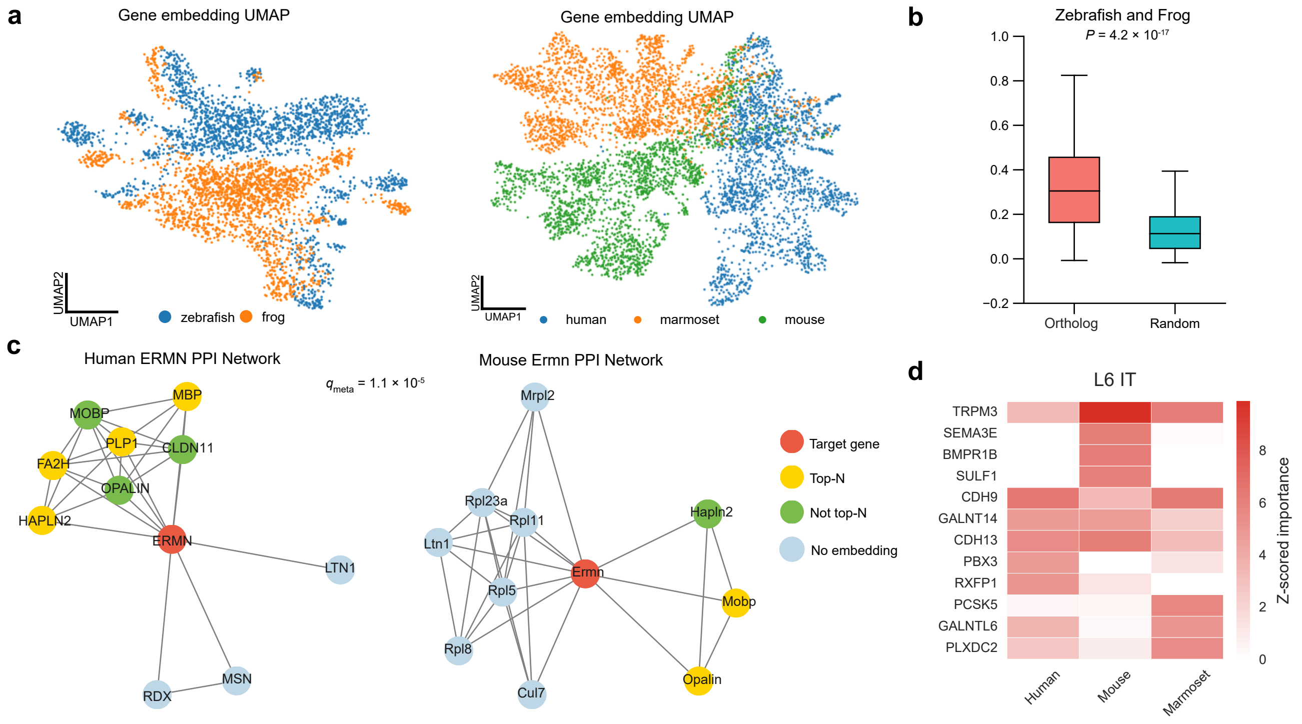


Supplementary Figure 11: CHORD learns aligned cross-species gene embeddings that capture biological relationships. (a) UMAP projection of the learned gene embedding for zebrafish and frog; each point is a gene, colored by species. (b) Distribution of cosine similarities in the gene embeddings for one-to-one ortholog pairs and an equal-sized set of randomly paired non-ortholog pairs between zebrafish and frog. (c) PPI networks for orthologs of *ERMN* in human and mouse. (d) Gene importance for L6 IT cell type across species (gradient×input). Heatmap shows the top $N = 5$ genes for each species (Human, Mouse, Marmoset), ranked by z-scored importance (within-species normalization).

Supplementary Table 1: Low-resolution evaluation groups for the primary motor cortex datasets and their corresponding high-resolution cell-type labels.

| High-res label | Low-res group | High-res label | Low-res group |
|---|---|---|---|
| Astro_1 | Astro | OPC | OPC |
| Astro_2 | Astro | Oligo_1 | Oligo |
| Chandelier | Chandelier | Oligo_2 | Oligo |
| Endo | Endo | Pvalb_1 | Pvalb |
| L2/3 IT | L2/3 IT | Pvalb_2 | Pvalb |
| L5 ET_1 | L5 ET | Sncg_1 | Sncg |
| L5 ET_2 | L5 ET | Sncg_2 | Sncg |
| L5 IT_1 | L5 IT | Sncg_3 | Sncg |
| L5 IT_2 | L5 IT | Sncg_4 | Sncg |
| L5 IT_3 | L5 IT | Sst Chodl | Sst Chodl |
| L5/6 NP | L5/6 NP | Sst_1 | Sst |
| L6 CT_1 | L6 CT | Sst_2 | Sst |
| L6 CT_2 | L6 CT | Sst_3 | Sst |
| L6 IT_1 | L6 IT | Sst_4 | Sst |
| L6 IT_2 | L6 IT | Sst_5 | Sst |
| L6 IT_3 | L6 IT | Sst_6 | Sst |
| L6b | L6b | Sst_7 | Sst |
| Lamp5_1 | Lamp5 | VLMC | VLMC |
| Lamp5_2 | Lamp5 | Vip_1 | Vip |
| Lamp5_3 | Lamp5 | Vip_2 | Vip |
| Lamp5_4 | Lamp5 | Vip_3 | Vip |
| Lamp5_5 | Lamp5 | Vip_4 | Vip |
| Microglia/PVM | Microglia/PVM | | |

Supplementary Table 2: Human–mouse ortholog pairs with PPI–embedding overlap: per-species $q$ values and meta-analysis $q_{\mathrm{meta}}$ ($q_{\mathrm{meta}} < 0.05$).

| Human gene | Mouse gene | $q_{\mathrm{human}}$ | $q_{\mathrm{mouse}}$ | $p_{\mathrm{meta}}$ | $q_{\mathrm{meta}}$ |
|---|---|---|---|---|---|
| ERMN | Ermn | $9.5 \times 10^{-4}$ | 0.11 | $2.0 \times 10^{-8}$ | $1.1 \times 10^{-5}$ |
| HEPACAM | Hepacam | 0.008 | 0.17 | $6.9 \times 10^{-7}$ | $2.0 \times 10^{-4}$ |
| AEBP1 | Aebp1 | 0.12 | 0.068 | $2.1 \times 10^{-6}$ | $4.1 \times 10^{-4}$ |
| IFI44 | Ifi44 | 0.027 | 0.78 | $1.2 \times 10^{-4}$ | 0.018 |
| PLP1 | Plp1 | 0.21 | 0.41 | $2.3 \times 10^{-4}$ | 0.026 |
| PARP14 | Parp14 | 0.008 | 1.00 | $3.4 \times 10^{-4}$ | 0.033 |
| GAD2 | Gad2 | 0.53 | 0.17 | $4.3 \times 10^{-4}$ | 0.036 |
